# ArtinM Cytotoxicity in B cells derived from Non-Hodgkin’s Lymphoma is regulated by CD45 phosphatase activity and Src family kinases

**DOI:** 10.1101/2022.07.05.498876

**Authors:** Bruno Rafael Barboza, Sandra Maria de Oliveira Thomaz, Airton de Carvalho, Enilza Maria Espreafico, Jackson Gabriel Miyamoto, Alexandre Keiji Tashima, Maurício Frota Camacho, André Zelanis, Maria Cristina Roque-Barreira, Thiago Aparecido da Silva

## Abstract

Receptors on the immune cell surface have a variety of glycans that may account for the immunomodulation induced by lectins, which have a carbohydrate recognition domain (CRD) that binds to monosaccharides or oligosaccharides in a specific manner. ArtinM, a D-mannose-binding lectin obtained from *Artocarpus heterophyllus*, has affinity for the N-glycans core. Immunomodulation by ArtinM toward the Th1 phenotype occurs via its interaction with TLR2/CD14 N-glycans on antigen-presenting cells, as well as recognition of CD3γ N-glycans on murine CD4+ and CD8+ T cells. ArtinM exerts a cytotoxic effect on Jurkat human leukemic T cell line and human myeloid leukemia cell line (NB4). The current study evaluated the effects of ArtinM on murine and human B cells derived from non-Hodgkin’s lymphoma. We found that murine B cells are recognized by ArtinM via the CRD, and the ArtinM stimulus did not augment the proliferation rate or production of IL-2. However, murine B cells incubation with ArtinM augmented the rate of apoptosis, and this cytotoxic effect of ArtinM was also seen in human B cell lines sourced from non-Hodgkin’s lymphoma Raji cell line. This cytotoxic effect was inhibited by the phosphatase activity of CD45 on Lck, and the protein kinases of the Src family contribute to cell death triggered by ArtinM.

## 1. Introduction

Glycosylation is a phenomenon in the post-translational modification of proteins [1]. The linkage of glycans (monosaccharides or oligosaccharides) to proteins and lipids gives rise to glycoconjugates [2]. The attached glycans may account for biological activities in different cell types [3], for example, innate and adaptive immune cells exhibit a panel of receptors for distinct classes of glycans [4–8] and their binding leads to cell activation and immunomodulation mediated by cytokine release and/or cell differentiation [9–12]. In addition, glycoproteins on the cell surface can mediate apoptosis of immune cells [13]. Glycans recognition by lectins may trigger biological responses [3]. Lectins are heterogeneous proteins whose carbohydrate recognition domain (CRD) react with mono-or oligosaccharides in a specific, reversible, and noncovalent manner. Studies on the interactions between lectins and glycoconjugates [14–16] have revealed their ability to modulate immune responses [17,18].

Our group previously reported the immunomodulatory activity of ArtinM, a D-mannose-binding lectin obtained from the seeds of *Artocarpus heterophyllus*. ArtinM is structurally organized as a homotetramer formed by 16-kDa non-glycosylated subunits, each polypeptide chain containing a CRD with an affinity for Manα1–3 [Manα1–6] Manβ1–4, which constitutes the N-glycans core [19–21]. Immunomodulation of the Th1 phenotype by ArtinM occurs via its interaction with TLR2/CD14 N-glycans on antigen-presenting cells, which induces IL-12 production [8,22]. In addition, ArtinM interacts with CD3γ N-glycans on murine CD4^+^ and CD8^+^ T cells, contributing to Th1 response and T cell proliferation [23]. ArtinM targets mast cells and neutrophils via interactions with FcεRI (high-affinity IgE receptor) [24] and CXCR2 (CXC-chemokine receptor type 2) [25], respectively. Recently, we demonstrated the capacity of ArtinM to induce murine B cell activation, detected by the augmented production of IL-17 and IL-12p40 cytokines, a response that does not depend on lectin binding to TLR2/CD14 [26]. In contrast, ArtinM exerts a cytotoxic effect on Jurkat human leukemic T cell line [23] and human myeloid leukemia cell line (NB4) [27]. These findings indicate that ArtinM induces distinct biological activities in several cell types, including tumor cells, via carbohydrate recognition.

B cells originate from hematopoietic stem cells (HSCs) in the bone marrow (BM) in adulthood [28]. HSC differentiation into mature B cells occurs at different stages, pro-B, pre-B, and immature B cells, characterized by the expression of cell surface-specific markers and successive rearrangement of the immunoglobulin (Ig) heavy (H) and light (L) gene segments [29]. B lymphocytes reach the final stage when Ig-kappa and/or Ig-lambda L chains rearrange over time during antigen-independent development [30]. The subdivision of B cells into B-1 and B-2 cells is based on their phenotype and functional properties. The two subpopulations are distinct in their ontogeny, anatomical distribution, and immune response [31–33]. B-1 cells show differential expression of CD5 and CD11b, which allows the characterization of B-1a and B-1b cells. B-1 cells are usually found in the peritoneal cavity [34] and are the leading producers of natural polyreactive antibodies [35]. The B-2 subpopulation comprises spleen marginal zone B cells (B-Mz) and follicular B cells (B-Fo) from the spleen and lymph node follicles. B-Mz cells, located at the interface between blood circulation and lymphoid tissue, have unique roles in innate immunity that account for the initiation of rapid natural antibodies production and efficient immune surveillance [36]. It is unlikely that B-Fo cells recirculate in the spleen and lymph node follicles and respond classically to antigens in a T-cell-dependent manner [37,38].

B-Fo cells are essential for structuring the germinal centers (GCs) of follicles, where antibody production, affinity maturation, and diversity occur [39]. Although GC reactions are strictly regulated, regulator genes are susceptible to mutations that favor malignant transformation of B cells and occurrence of lymphomas [40]. Most lymphomas resulting from GC reactions are non-Hodgkin’s lymphoma (B-NHL) types [41,42]. Burkitt lymphoma is a highly aggressive B-NHL subtype [43]. The molecular signature of all BL subtypes involves chromosomal rearrangement with translocation of the c-MYC gene (8; 14) into the immunoglobulin heavy chain locus [44,45]. Several other genes have been reported as potential inductors of BL tumorigenesis, such as those involved in the BCR, PI3K/AKT signaling pathways, apoptosis, epigenetic regulation, and G protein-coupled receptors [46].

In this study, we report new molecular mechanisms used by ArtinM to induce apoptosis in B-NHL cells. Herein, we demonstrated that ArtinM at low concentrations increased the mitochondrial activity of murine B cells without causing cell proliferation. We also verified apoptosis in murine B cells and the cytotoxic effect of ArtinM in human B cells derived from NHL. Carbohydrate recognition accounts for Raji and Daudi cell apoptosis induced by ArtinM in a manner dependent on protein kinases of the Src family and phosphatase activity of CD45. Quantitative proteomics revealed low levels of proteins associated with cell proliferation and survival regulation in ArtinM-stimulated Raji cells. Our findings provide new directions to be considered in delineating strategies for cancer therapy based on carbohydrate recognition.

## 2. Results

### 2.1. ArtinM increases the mitochondrial activity of B cells without inducing cell proliferation

The immunomodulatory activity of ArtinM on antigen-presenting cells and T cells has been well studied, whereas the effect of ArtinM on B lymphocytes has not been sufficiently explored. In this study, we first assessed the effects of ArtinM on purified murine splenic B cells. Mitochondrial activity was measured using the MTT assay 24 and 48 h after incubation with ArtinM (0.312–5 µg/mL) (Figure 1A). Mitochondrial activity did not increase after 24 h incubation. However, at 48 h, stimulation with 0.312 and 0.625 µg/mL ArtinM increased the mitochondrial activity of B cells by 28.5% and 18.7%, respectively, in comparison to unstimulated cells (Figure 1A). Because augmentation of mitochondrial activity is frequently related to cell proliferation, the 48 h ArtinM-stimulated cells were evaluated using the tritiated thymidine ([^3^H]-TdR) incorporation assay, and IL-2 production was measured using enzyme-linked immunosorbent assay (ELISA). Compared with unstimulated cells (medium), B cells stimulated with any ArtinM concentration (0.312–5 µg/mL) did not show increased [^3^H]-TdR incorporation (Figure 1B), increase in the proliferation rate stimulation index (Figure 1C), or IL-2 levels (Figure 1D). Therefore, ArtinM stimulated the mitochondrial activity of murine B cells without inducing cell proliferation.

**Figure 1.**
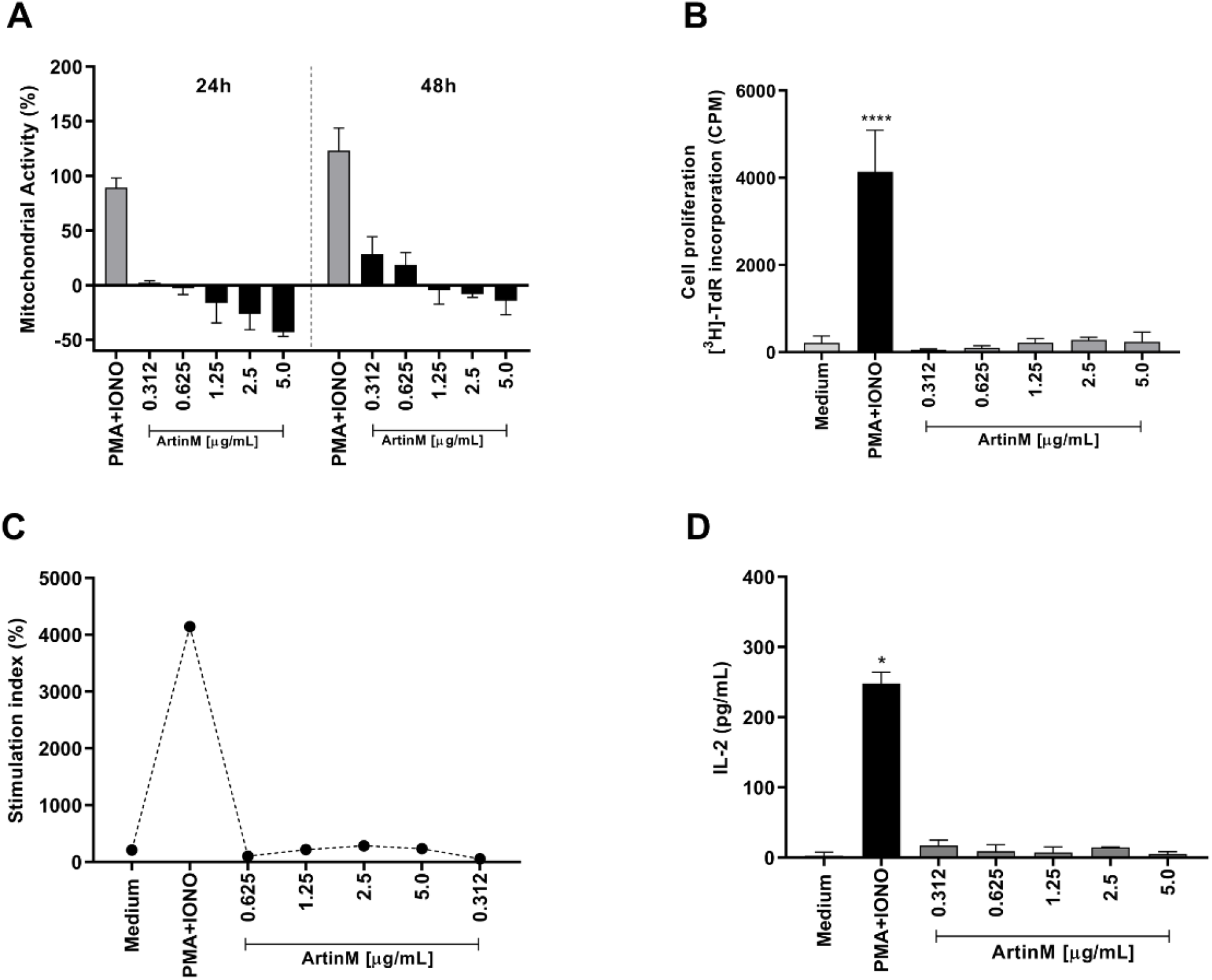
Increased mitochondrial activity of murine B cells induced by ArtinM is not accompanied by B cell proliferation or IL-2 production. B cells were purified from a spleen cell suspension obtained from C57BL/6 mice, seeded in 96-well microplates (5 × 10^5^ cells/well), and incubated with ArtinM at different concentrations (**A-D**). A mixture of Phorbol-12-myristate-13-acetate (PMA; 50 ng/mL) and Ionomycin (IONO; 1 µM) was used as a positive control for cell activation, and medium alone was used as a negative control (Medium). (**A**) Following 24 or 48 h incubation with ArtinM, 3-(4,5-dimethyl-2-thiazolyl)-2,5-diphenyl-2H-tetrazolium bromide (MTT; 50 µg/mL) was added to the cells, and mitochondrial activity was estimated by MTT reduction and expressed as a percentage calculated from the ratio between the absorbance of stimulated and non-stimulated B cells. (**B, C**) Following 48 h incubation with ArtinM, the proliferative rate of murine B cells was calculated according to thymidine incorporation ([^3^H]-TdR) (0.5µCi/well); results are expressed as (B) count per minute (CPM) and (**C**) stimulation index of cell proliferation. (**D**) IL-2 levels in the supernatant of ArtinM-stimulated B cells, for 48 h, were measured using ELISA; the values are expressed in pg/mL. Significant differences compared to the negative control are shown by *, *p* < 0.05 and ***, *p* < 0.0001.

### 2.2. ArtinM induces apoptosis of murine B cells

Considering our observation of reduced mitochondrial activity in murine B cells incubated for 24 h with ArtinM at high concentrations (Figure 1A), we examined the occurrence of apoptosis in ArtinM-stimulated B cells. Following 24 h incubation with 0.312–5.0 μg/mL ArtinM, we assayed B cells for annexin V (AnV) and propidium iodide (PI) markers. Compared with unstimulated B cells, B cells incubated with high concentrations of ArtinM showed increase in AnV and PI (AnV+/PI+) double-positive cells in a dose-dependent manner (Figure 2A and 2B), indicating that ArtinM induces apoptosis in murine B cells. Then, the immunomodulatory activity of ArtinM at low concentrations in B cells was examined by verifying the relative frequency of IL-12-and IFN-γ-positive cells. Compared to unstimulated cells, stimulation with ArtinM at low concentrations (0.312–1.25 µg/mL) did not increase the frequency of IL-12+/IFN-γ+ double-positive B cells (Figure 2C). These data highlight the ArtinM preference for inducing apoptosis in murine B cells over immunomodulation for Th1 cytokine production.

**Figure 2.**
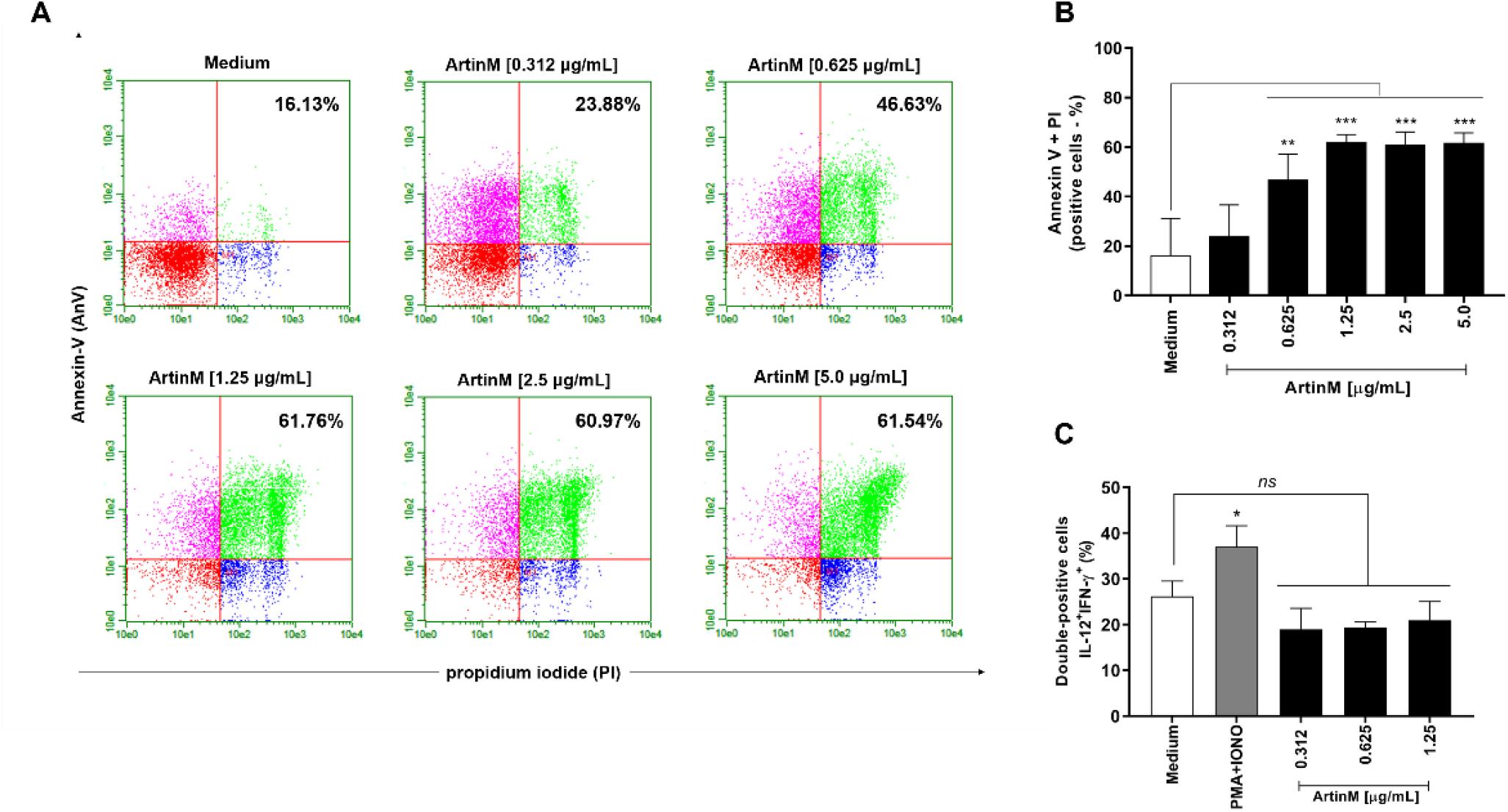
ArtinM increases the frequency of annexin V- and PI-positive B cells. B cells purified from a suspension of C57BL/6 mice spleen cells were seeded in 96-well microplates (5 × 10^5^ cells/well) and incubated for 24 h with ArtinM at several concentrations. Medium alone was used as a negative control (Medium). (**A, B**) B cells were labeled with annexin V FITC and propidium iodide (PI) to determine the frequency of apoptotic cells (annexin V + PI) by flow cytometry. (**C**) The frequency of IL-12+/IFN-γ+ double-positive cells was also determined by flow cytometry. A mixture of phorbol-12-myristate-13-acetate (PMA; 50 ng/mL) plus Ionomycin (IONO; 1 µM) was used as a positive control. Results are expressed as mean ± standard error of the mean (SEM), and three independent assays were performed. Significant differences compared to the negative control are shown by *, *p* < 0.05 and ***, *p* < 0.0001. ns: not significant.

### 2.3. ArtinM interacts with Raji and Daudi cells via its CRD but induces only Raji cells apoptosis

After verifying that ArtinM induces apoptosis of murine B cells, we investigated the cytotoxic effect of lectin on human B cell lines sourced from NHL, i.e., Raji and Daudi cells. First, we verified the binding of ArtinM to the surface of both the cell lines (Figure 3A and 3B). In addition, we evaluated the inhibition of ArtinM binding by specific (mannose or mannotriose) and non-specific (lactose) sugars. As expected, only the specific sugars inhibited ArtinM binding to the cell surface. We then assayed propidium iodide incorporation by each cell line in response to ArtinM (0.02–20 µg/mL). The data obtained for Raji cells allowed us to verify the effect of lectin on cell growth and viability compared to the negative control (medium). The reduction in cell growth and viability in Raji cells was dose-dependent from 0.3 to 5 µg/mL ArtinM. In contrast, the growth and viability of Daudi cells were not affected by ArtinM at concentrations ranging from 0.625 µg/mL to 10 µg/mL. The cytotoxic effect of ArtinM on B-NHL cell lines was also demonstrated by determining the frequency of Raji and Daudi cells for annexin V and propidium iodide positivity (AnV+/PI+) following 48 h-stimulation with ArtinM. A dose-dependent increase in the frequency of double-positive Raji cells was observed (Figure 4A and 4B). In contrast, the percentage of annexin V- and PI-positive Daudi cells did not increase with ArtinM 48 h-stimulation (Figure 4, panels A and C). These findings indicate that ArtinM exerts its cytotoxic effect on Raji cells via apoptosis.

**Figure 3.**
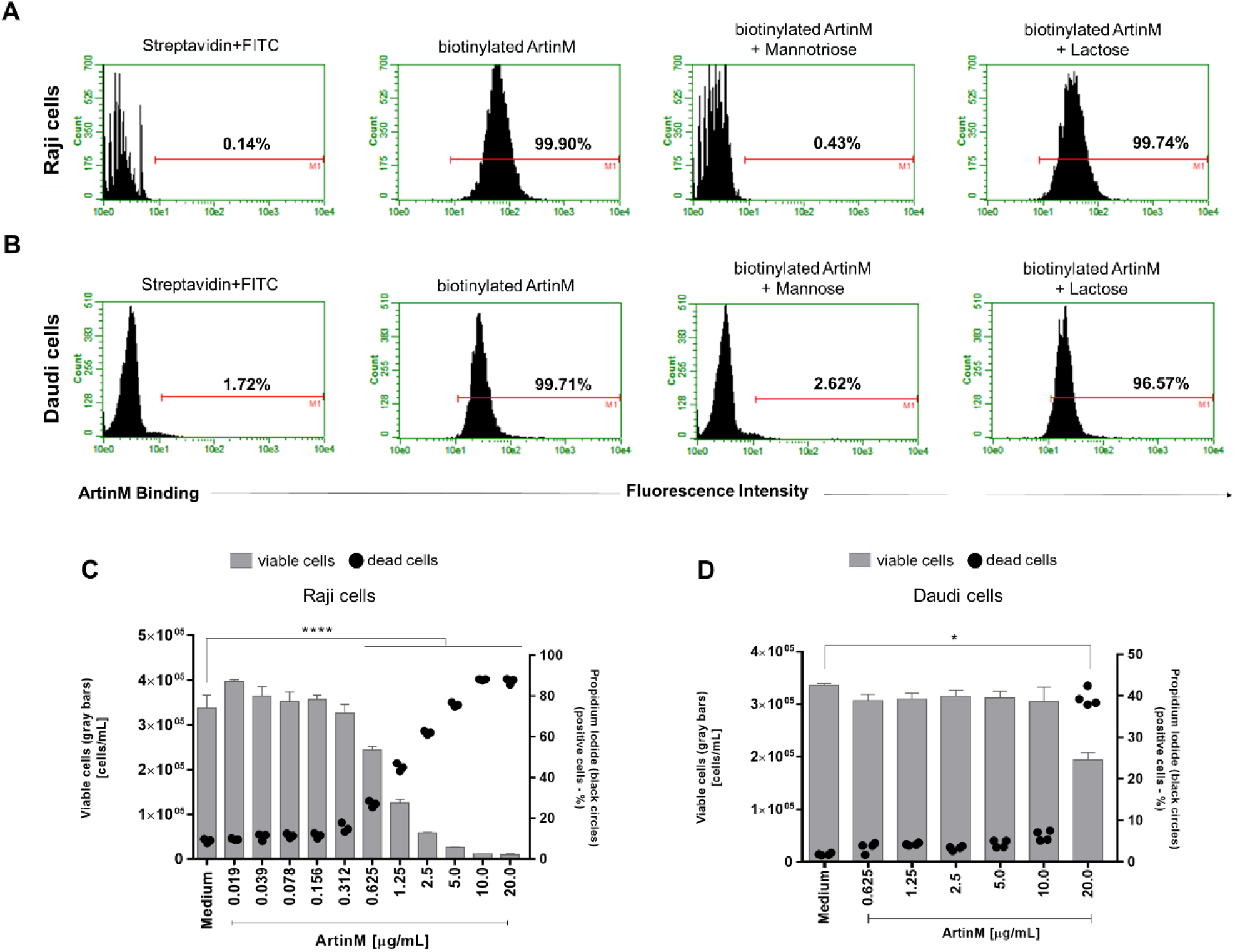
ArtinM binds Raji and Daudi cell surface through carbohydrate recognition, reducing Raji cells’ viability and increasing death. (**A**) Raji and (**B**) Daudi cells (1 × 10^6^ cells/mL) were fixed and incubated with biotinylated ArtinM (20 µg/mL), pre-treated or not with mannotriose (1 mM), Mannose (20 mM) or lactose (20 mM). After washing, streptavidin-FITC (5 µg/mL) was added to the cells, which were analyzed by flow cytometry. Streptavidin-FITC alone was used as a negative control. Histograms represent the fluorescence intensity of cells for ArtinM detection; the positive range is indicated by the red line (M1). (**C**) Raji and (**D**) Daudi cells (1 × 10^5^ cells/mL) were distributed in 96-well microplates and incubated for 48 h at 37 °C with ArtinM at different concentrations. Medium alone was used as a negative control (Medium). The percentage of live and the frequency of dead (propidium iodide(PI)-positive) cells was determined by flow cytometry. The number of viable cells (grey bars) and the frequency of PI-positive cells (black circles) are shown in C and D. The results are expressed as mean ± standard error of the mean (SEM); data are representative of three experimental replicates. Significant differences compared to the negative control are shown by *, *p* < 0.05 and ***, *p* < 0.0001.

**Figure 4.**
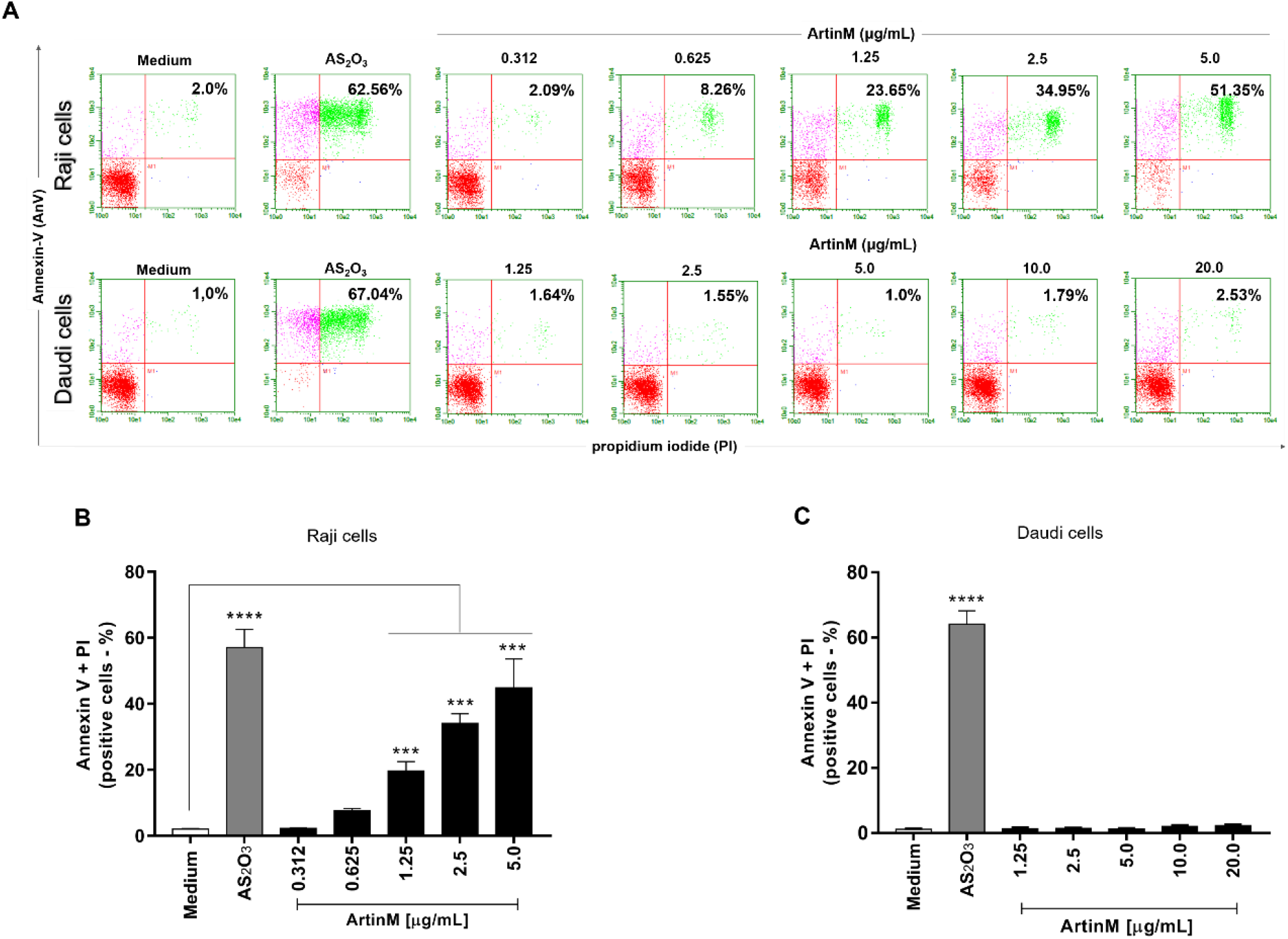
ArtinM induces apoptosis of Raji cells. Raji and Daudi cells (1 × 10^5^ cells/mL) were seeded in 96-well microplates and incubated for 48 h at 37 °C with 0.312 to 5.0 µg/mL and 1.25 to 20.0 µg/mL ArtinM, respectively. Arsenic trioxide (AS_2_O_3_) at concentrations of 12 µM (Raji cells) and 24 µM (Daudi cells) was used as a positive control for induction of cell death and medium alone was used as a negative control (Medium). The cells were incubated with annexin V-FITC (5.0 μg/mL) and propidium iodide (10.0 μg/mL), and the frequency of double-labeled (AnV+/PI+) Raji (**A, B**) and Daudi (**C**) cells was determined by flow cytometry. Values were expressed as mean ± standard error of the mean (SEM). Significant differences compared to the negative control are shown by ***, *p* < 0.001 and ****, *p* < 0.0001.

### 2.4. Raji and Daudi cells did not undergo DNA fragmentation or increased expression of mitochondrial and autophagy markers in response to ArtinM

The potent cytotoxic effects of ArtinM on Raji cells led us to investigate DNA damage in Raji and Daudi cells incubated with ArtinM for 24 and 48 h. Agarose gel electrophoresis revealed that the ArtinM-stimulated cells did not show DNA cleavage, whereas treatment with AS_2_O_3_ led to DNA smearing (Figure 5A). Moreover, 48h-stimulated Raji cells did not show increased apoptosis markers associated with the mitochondrial pathway (Caspase-3, Apaf-1, and Smac/DIABLO) or autophagy (LC3-I, ATG14, and ATG12), as measured by the relative expression of their transcripts by qRT-PCR (Figure 5B-G). Considering the pronounced cytotoxic effect of ArtinM on Raji cells and the stable relative expression of all studied markers in response to ArtinM compared to unstimulated cells, we infer that the apoptosis of Raji cells induced by ArtinM is not related to the mitochondrial pathway or autophagy.

**Figure 5.**
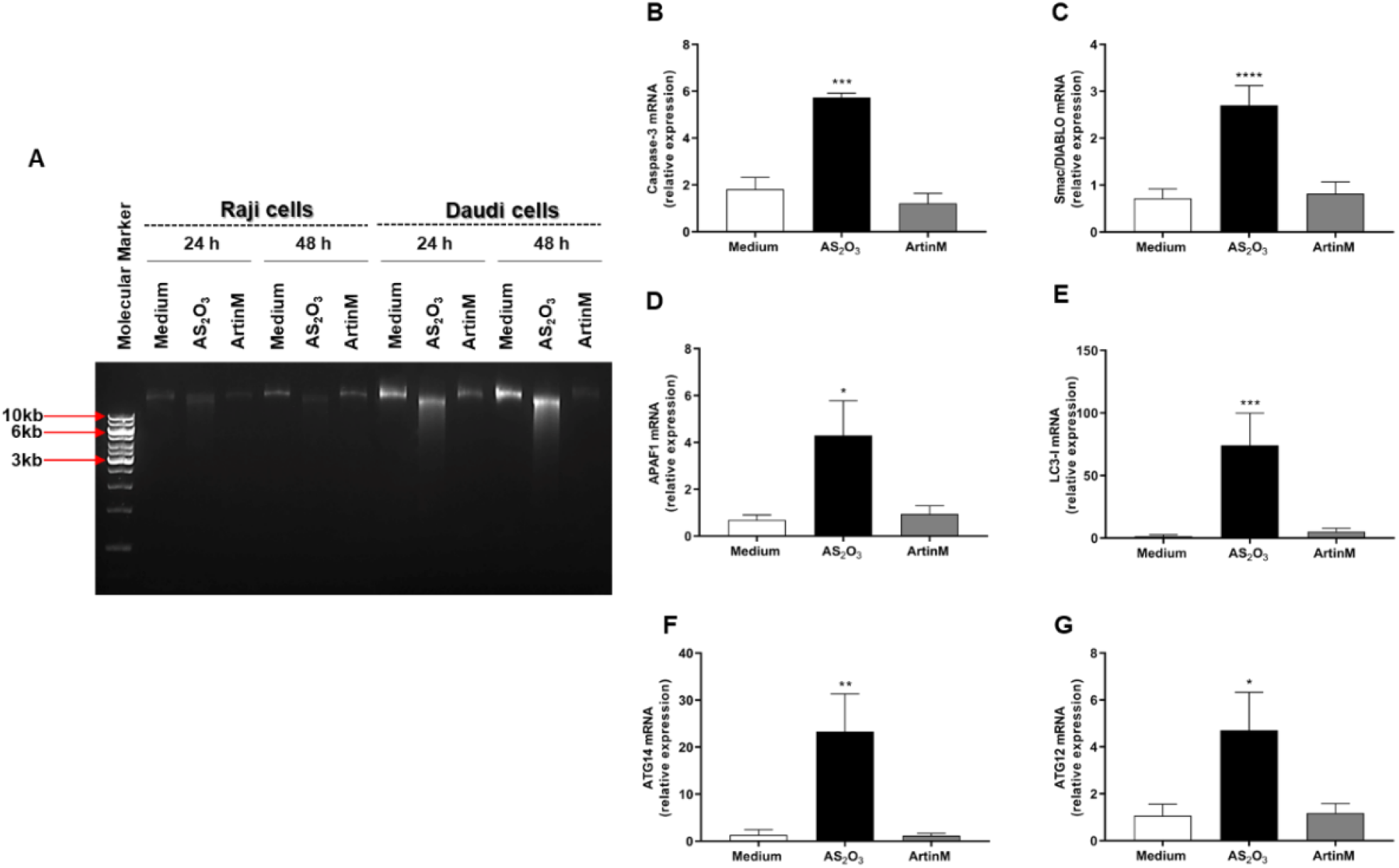
Apoptosis of ArtinM-stimulated Raji cells is not associated with DNA fragmentation and does not involve the mitochondrial or autophagy pathways. Raji or Daudi cells (1 × 10^5^ cells/mL) were seeded in 96-well microplates and stimulated with ArtinM at concentrations of 1.25 µg/mL (Raji cells) or 20.0 µg/mL (Daudi cells) for 24 or 48 h at 37 °C. Arsenic trioxide (AS_2_O_3_) at concentrations of 12 µM (Raji cells) and 24 µM (Daudi cells) was used as a positive control of cell death induction; culture medium alone (Medium) was a negative control. (**A**) DNA extracts were visualized by agarose gel electrophoresis (0.8%) after staining with SYBR safe. (B-G) The relative expression of Caspase-3 (**B**), Smac/DIABLO (**C**), APAF1 (**D**), LC3-I (**E**), ATG14 (**F**), and ATG12 (**G**) was measured by qRT-PCR in ArtinM-stimulated Raji cells for 48 h. The Ct values of the target transcripts were normalized to the relative expression of β-actin as an endogenous control. The results are expressed as mean ± SEM, and the differences were considered significant at *p* < 0.05 (*), *p* < 0.001 (**), *p* < 0.001 (***), or *p* < 0.0001 (****).

### 2.5. ArtinM-induced Raji cell apoptosis is not affected by by pharmacological inhibitors of the p38 MAPK, JNK, ERK, PKC, and PTK signalling molecules

Because the effect of ArtinM on apoptosis induction was confirmed in Raji cells (Figures 3 and 4), we utilized the Raji cell line to search for specific signaling pathways that account for lectin-stimulated apoptosis. Next, we examined whether the apoptotic response to ArtinM was maintained in Raji cells pre-treated with pharmacological inhibitors of p38MAPK (SB202190), JNK (SP600125), ERK (PD98059), PKC (H-7), and PTK (genistein) signaling molecules. Raji cells were incubated with a pharmacological inhibitor (20 µM) for 210 min, followed by stimulation with ArtinM (1.25 µg/mL). After 24 h, flow cytometry (Figure 6) was performed, it was shown that the pharmacological inhibitors did not affect the number of viable cells (light gray bars, Figure 6A) or the frequency of apoptotic cells provided by the negative control (medium) (Figure 6B and 6C). In addition, the inhibitors, except genistein, did not affect the lower viable cell counts by the ArtinM stimulus. Genistein, a PTK inhibitor, provided a cell count that was lower than that determined using ArtinM alone (dark grey bars, Figure 6A). We also analyzed the relative frequency of double-positive cells (AnV+/PI+) by flow cytometry. Cells pretreated with the inhibitors and ArtinM-stimulated cells contained a percentage of AnV+/PI+ cells that did not differ from those stimulated with ArtinM alone (Figure 6B and 6C). Although the PTK inhibitor reduced cell viability, it did not affect the apoptosis of Raji cells induced by ArtinM. Therefore, the effect of ArtinM on Raji cells was maintained even after selective pharmacological blocking of the signaling molecules p38, JNK, ERK, PKC, or PTK, indicating that none of the assayed signaling molecules accounted for ArtinM-induced apoptosis of Raji cells.

**Figure 6.**
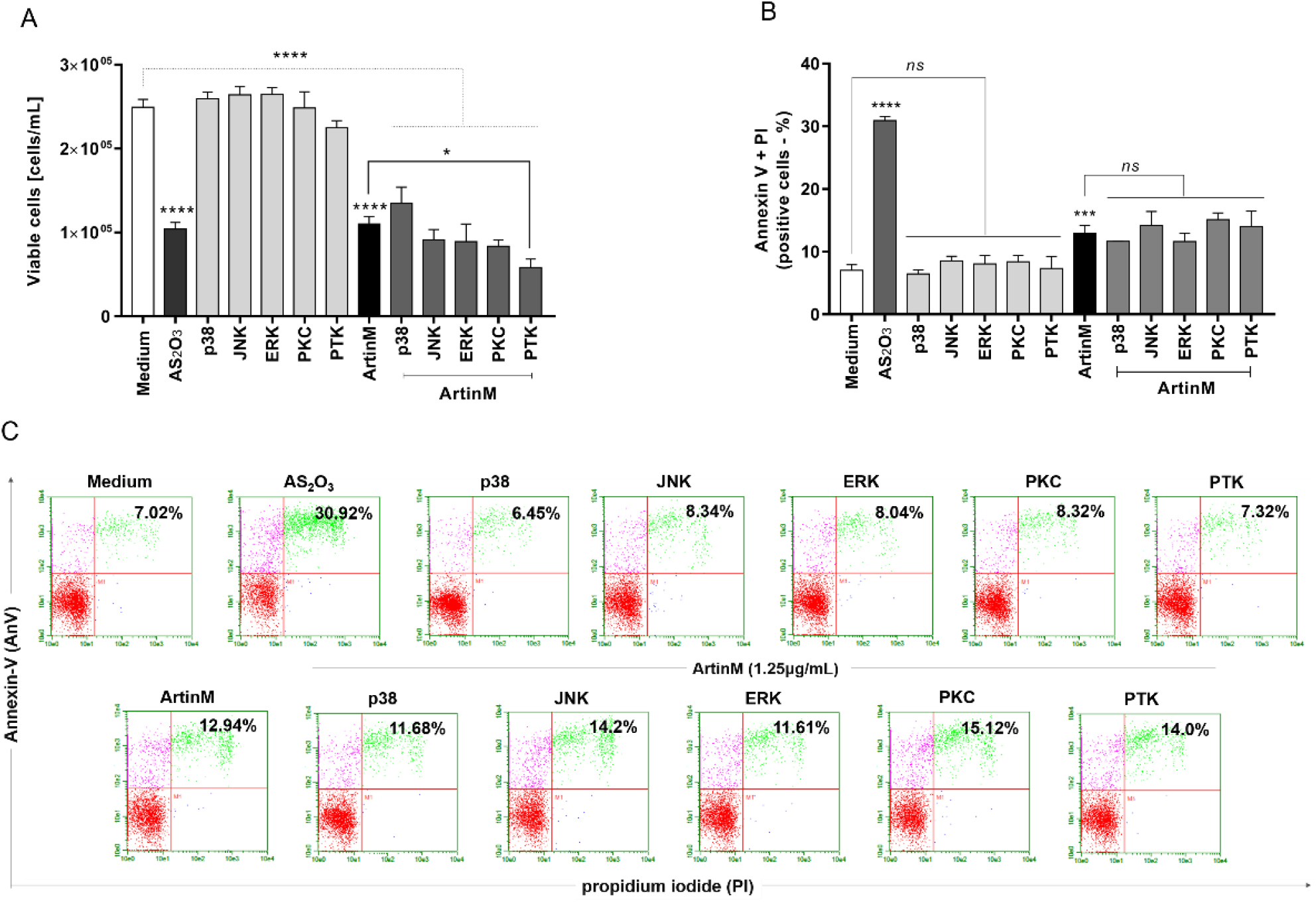
Pharmacological inhibition of the signaling molecules p38, JNK, ERK, PKC, or PTK does not affect ArtinM-induced apoptosis of Raji cells. Raji cells (1 × 10^5^ cells/mL) were seeded in 96-well microplates and pre-treated or not with pharmacological inhibitors (20µM), such as SB202190 (p38MAPK inhibitor), SP600125 (JNK inhibitor), PD98059 (ERK inhibitor), H-7 (PKC inhibitor) and genistein (PTK inhibitor). After 210 min, the cells were stimulated with ArtinM (1.25 µg/mL) for 24 h at 37 °C. Arsenic trioxide (AS_2_O_3_; 12 μM) was used as a positive control of cell death induction, and the medium alone was used as a negative control (Medium). (**A**) The number of viable cells (cells/mL) was determined as the difference between the total number of cells and PI+ cells (dead cells), detected by flow cytometry. The results are expressed as mean ± standard error of three independent experiments in triplicates. (**B, C**) The relative frequency of apoptotic cells was determined by annexin V-FITC (5.0 μg/mL) and PI (10.0 μg/mL) staining; double-positive cells (AnV+/PI+) were detected by flow cytometry analysis, and their relative frequency was represented as a percentage. Values with significant differences in relation to the negative control (medium) or cells stimulated only with ArtinM are shown by *, *p* < 0.05; ***, *p* < 0.001; ****, *p* <0.0001. ns: not significant.

### 2.6. Phosphatase activity of CD45 and protein kinases of the Src family account for the cytotoxic effect of ArtinM on Raji cells

To further investigate the signaling pathways responsible for the cytotoxic effect of ArtinM on Raji cells, we evaluated whether inhibition of protein tyrosine kinases could modify apoptosis induced by ArtinM. We chose to study the effect of phosphatase activity of the CD45 receptor and protein kinases of the Src family (Lyn, Fyn, and Blk) because of their known roles in regulating cell survival. Raji and Daudi cells were pre-treated with NQ-301, an inhibitor of dephosphorylation of the Y505 residue of the Lck molecule (lymphocyte-specific protein tyrosine kinase) of the CD45 receptor, or with Dasatinib, an inhibitor of the activity of Lyn, Fyn, and Blk kinases. The inhibitors were used at different concentrations (0.1, 0.2, and 0.5 μM) and, after 210 min, the cells were stimulated or not with ArtinM (1.25 µg/mL for Raji cells; 20.0 µg/mL for Daudi cells) for 24 h. We deduced the number of viable cells per mL by subtracting the number of PI+ cells detected by flow cytometry from the total number of cells. In Raji cells, treatment with either inhibitor (NQ-301 or dasatinib), not followed by stimulation with ArtinM, had no effect on cell viability (Figure 7A and 7E, light grey bars compared to the white bar). The same treatments modified the effect of ArtinM on the number of viable Raji cells, which was significantly reduced by NQ-301 (Figure 7A, dark gray bars compared to the black bar) and increased by dasatinib (Figure 7E, dark gray bars compared to the black bar). Conversely, the percentage of Raji cells that had incorporated PI, increased or decreased when stimulation with ArtinM was preceded by NQ-301 or dasatinib treatment, respectively (Figure 7B and 7F).

**Figure 7.**
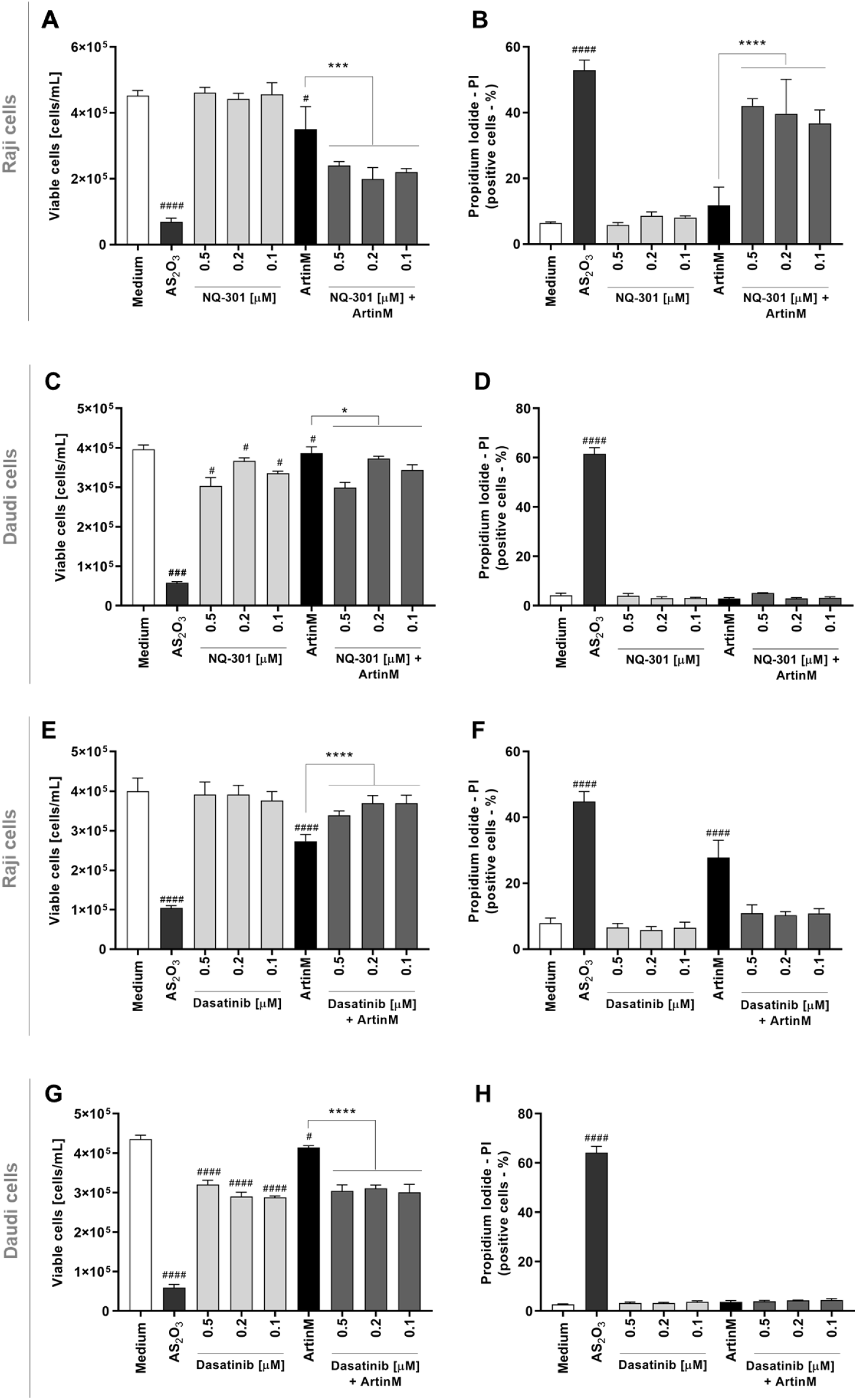
Pharmacological inhibitors of CD45 phosphatase activity and protein tyrosine kinases of the SCR family modify the cytotoxic effect of ArtinM in Raji cells. Raji (A-B and E-F) and Daudi (C-D, and G-H) cells (1 × 10^5^ cells/mL), seeded in 96-well microplates, were pre-treated or not with the pharmacological inhibitors NQ-301 (**A-D**) or dasatinib (**E-H**) at different concentrations (0.5 to 0.1 µM) for 210 minutes, followed by stimulation with 1.25 µg/mL ArtinM (Raji cells) or 20 µg/mL ArtinM (Daudi cells). Arsenic trioxide (AS_2_O_3_) at 12 µM was used in Raji cells, and at 24 µM in Daudi cells, as a positive control. The medium was used as a negative control. After 24 h, propidium iodide (10 μg/mL) was added to the cell culture and its incorporation was detected by flow cytometry, allowing the quantification of viable and non-viable cells. Based on the total cell number and the frequency of PI+ cells (dead cells, black bars), the quantification of viable cells (white bars) was determined. The results are expressed as mean ± standard error of the mean (SEM) of three independent experiments carried out in triplicates. Significant differences compared to the negative control (Medium) are shown by ^#^, *p* < 0.05; ^###^, *p* < 0.001; ^####^, *p* < 0.0001. Significant differences compared to the ArtinM alone are shown by *, *p* < 0.05; ***, *p* < 0.001; ****, *p* < 0.0001.

Only AS_2_O_3_ (positive control) promoted death in Daudi cells (Figure 7C-D and 7G-H). Upon stimulation with ArtinM or treatment with either inhibitor, the relative frequency of dead Daudi cells (PI+) remained as low as that observed in the negative control (medium) (Figure 7D and 7H). Treatment with either inhibitor, not followed by stimulation with ArtinM, significantly diminished the number of viable Daudi cells whereas (Figure 7D). Increased viability and decreased apoptosis in Raji cells following pre-treatment with dasatinib in response to ArtinM (Figure 7C) indicated that Lyn, Fyn, and Blk kinases play a role in the cytotoxic effect of ArtinM on Raji cells. This result evidenced that dasatinib exerted its activity only on ArtinM-stimulated Raji cells and not on unstimulated Raji cells (Figure 7G). Our data indicated that the phosphatase activity of CD45 on Lck inhibits the cytotoxic effect of ArtinM in Raji cells. In contrast, protein kinases of the Src family favor cell death triggered by ArtinM.

To validate that NQ-301 and dasatinib regulate cell death triggered by ArtinM, we used Raji cells pretreated with these inhibitors. We stimulated Raji cells with ArtinM and analyzed them for apoptosis markers to determine the frequency of AnV+ (early apoptosis marker) and AnV+/PI+ (late apoptosis marker) cells. We found that both pre-treatments with NQ-301 and dasatinib led to a reduced frequency of AnV+/PI+ Raji cells in response to ArtinM, compared to cells stimulated with ArtinM alone (Figure 8C and 8E). However, AnV+ Raji cell frequency in response to ArtinM was augmented by NQ-301 (Figure 8B) and was not altered by dasatinib pre-treatment (Figure 8D). These data indicate the negative regulation by CD45 in ArtinM-stimulated apoptosis of Raji cells, while the mediation of the cytotoxic effect of ArtinM by protein kinases of the Src family can be seen in Raji cells. To evaluate the effect of NQ-301 and dasatinib on ArtinM-stimulated apoptosis in normal B cells, they were purified from a suspension of mouse spleen cells. We treated the B cells with NQ-301 (0.1 μM) or dasatinib (0.1 μM), followed by stimulation with ArtinM (1.25 μg/mL). Pre-treatments only did not induce an apoptotic response in B cells, and ArtinM stimulus increased the frequency of AnV+/PI+ cells as expected. We detected a higher frequency of AnV+/PI+ cells in ArtinM-stimulated B cells pretreated with NQ-301, but not in dasatinib-treated cells (Figure 9). Therefore, Src family protein kinases, and not CD45, participate in ArtinM-induced apoptosis of murine B cells.

**Figure 8.**
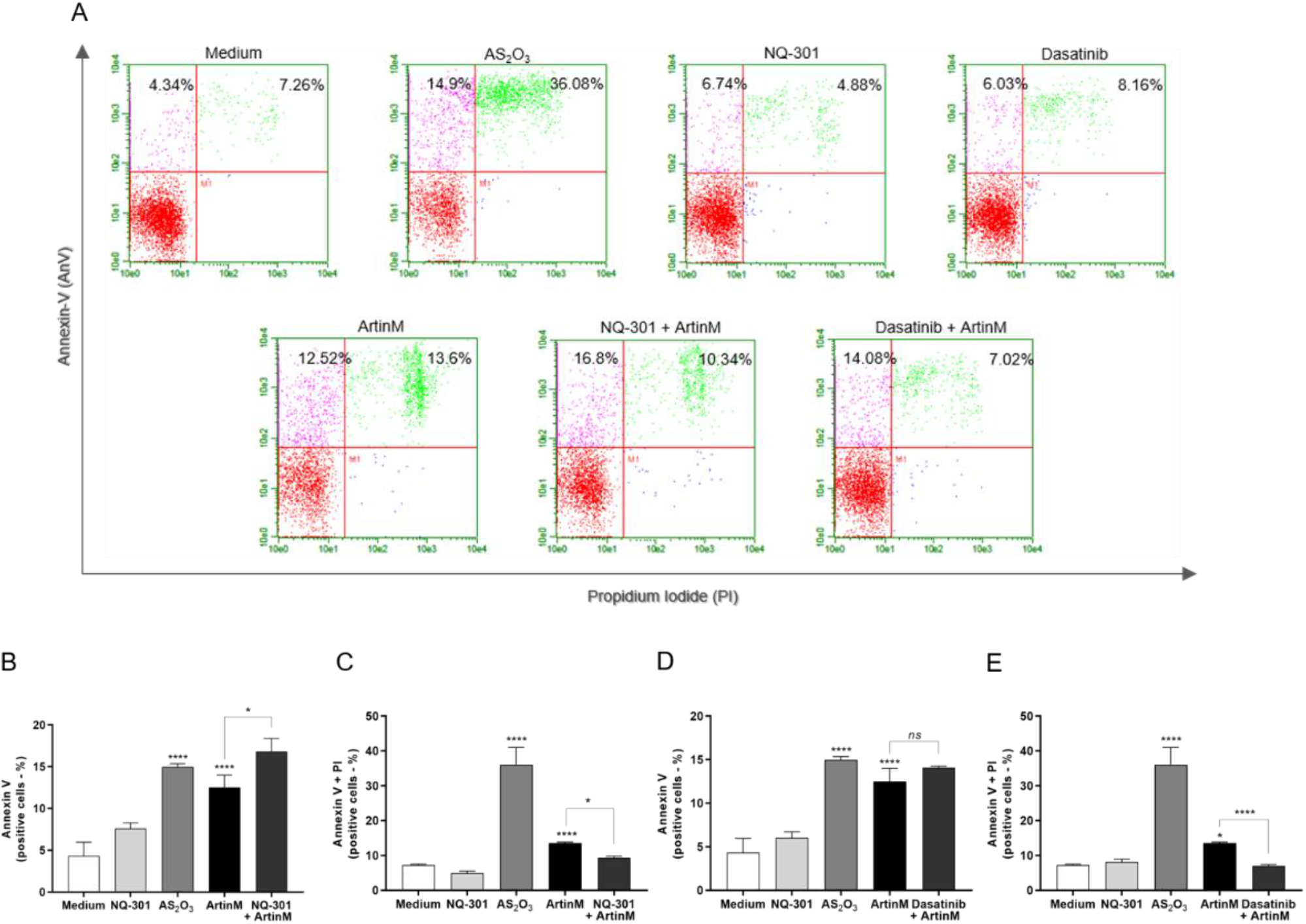
NQ-301 and dasatinib differently affect apoptosis of ArtinM-stimulated Raji cells. Raji cells (1 × 10^5^ cells/mL) were seeded in 96-well microplates and pre-treated or not with the pharmacological inhibitors NQ-301 **(A, B, and C)** or dasatinib **(A, D, and E)** at a concentration of 0.2 µM for 210 minutes. The cells were stimulated with ArtinM (1.25 µg/mL). Arsenic trioxide (AS_2_O_3_; 12 µM) was used as a positive control of cell death, and medium alone as a negative control. After 24 h stimulation, the cells were incubated with AnV-FITC (5.0 μg/mL) and PI (10.0 μg/mL) and analyzed by flow cytometry to determine the relative frequency (%) of Raji cells labeled only with annexin V (AnV+) **(B and D)** or double-labeled annexin V-FITC plus PI (AnV+/PI+) **(C and E)**. Significant differences compared to the negative control (Medium) or the isolated stimulus with ArtinM are shown by *, *p* < 0.05; ****, *p* < 0. 0001.

**Figure 9.**
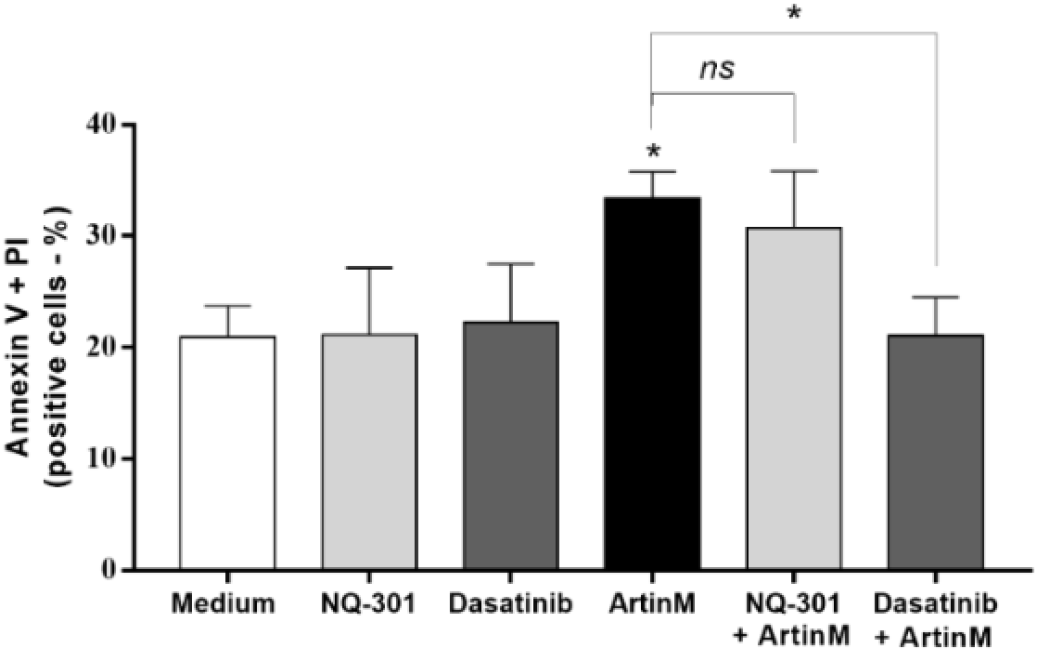
Dasatinib inhibits apoptosis of murine B cells induced by ArtinM. Murine B cells (5 × 10^5^ cells/mL) seeded in 96-well microplates were pre-treated or not with 0.1 µM NQ-301 or dasatinib for 210 minutes. The cells were stimulated or not with ArtinM (0.625 µg/mL). The medium was used as a negative control. After 24 h, the cells were labeled with annexin V-FITC (5.0 μg/mL) and PI (10.0 μg/mL) and the frequency (%) of AnV+/PI+ B cells was determined by flow cytometry. Significant differences compared to the negative control (medium) or stimulus with ArtinM are shown by *, *p* < 0.05.

### 2.7. Quantitative proteomics analysis to identify possible targets whose expression is regulated by the cytotoxic effect of ArtinM

A quantitative proteomics approach was used to identify cell signaling molecules possibly involved in the regulation of the survival of Raji and Daudi cells stimulated with ArtinM. We searched for proteins whose differential expression in NHL B cell lines was associated with the cytotoxic effect of ArtinM. A total of 2,091 proteins were identified, of which, 921 were quantified for both cell lines under different experimental conditions (Table 1A). Table 1 lists the top 10 upregulated and downregulated proteins in both Raji and Daudi cells following stimulation with ArtinM. In Raji cells, we identified upregulated proteins (Table 1A), including HLA-B (P01889) and HLA-DRA (P01903), involved in B-cell death. Eukaryotic translation initiation factors, such as eIF3-E (P60228) and eIF3-M (Q7L2H7), were also upregulated. Downregulated proteins, including those associated with the regulation of cell proliferation and survival, such as MIF (P14174) and PDLIM1 (O00151), were identified in ArtinM-stimulated Raji cells (Table 1B). In Daudi cells (Table 1C), positive regulators of cell cycle progression, namely, ANP32B (Q92688), GSPT1 (P15170), and SUB1 (P53999) were upregulated after ArtinM stimulus. In addition, ArtinM-stimulation of Daudi cells up-regulated proteins was involved in the transcriptional regulation of histones H1-2 (P16403), H1-4 (P10412), and H2AC11 (P0C0S8). In contrast, essential proteins that inhibit cell proliferation and induce apoptosis, such as NAT10 (Q9H0A0), PSMB2 (P49721), and RAB7A (P51149), were downregulated in the ArtinM-stimulated Daudi cells (Table 1D).

**Table 1.** The top 10 upregulated and downregulated proteins in Raji (A-B) and Daudi (C-D) cells after treatment with ArtinM.

| (A) |  |  |  |
| --- | --- | --- | --- |
| Gene name | Protein name | Accession | Fold-change |
| HLA-B | HLA class I histocompatibility antigen, B alpha chain | P01889 | -1.02275 |
| IPO7 | Importin-7 | O95373 | -0.77898 |
| HLA-DRA | HLA class II histocompatibility antigen, DR alpha chain | P01903 | -0.71939 |
| eIF3-E | Eukaryotic translation initiation factor 3 subunit E | P60228 | -0.67789 |
| STAT5A | Signal transducer and activator of transcription 5A | P42229 | -0.60567 |
| RPL30 | 60S ribosomal protein L30 | P62888 | -0.58276 |
| SURF4 | Surfeit locus protein 4 | O15260 | -0.55646 |
| eIF3-M | Eukaryotic translation initiation factor 3 subunit M | Q7L2H7 | -0.55029 |
| ARF3 | ADP-ribosylation factor 3 | P61204 | -0.52062 |
| NME1 | Nucleoside diphosphate kinase A | P15531 | -0.51406 |
| (B) |  |  |  |
| Gene name | Protein name | Accession | Fold-change |
| MIF | Macrophage migration inhibitory factor | P14174 | 0.491055 |
| IFITM1 | Interferon-induced transmembrane protein 1 | P13164 | 0.61364 |
| IFI30 | Gamma-interferon-inducible lysosomal thiol reductase | P13284 | 0.628365 |
| PDLIM1 | PDZ and LIM domain protein 1 | O00151 | 0.730311 |
| H2AC4 | Histone H2A type 1-B/E | P04908 | 0.790737 |
| HSPA4L | Heat shock 70 kDa protein 4L | O95757 | 0.804528 |
| BANF1 | Barrier-to-autointegration factor | O75531 | 0.845222 |
| H1-2 | Histone H1.2 | P16403 | 0.847602 |
| DENND4C | DENN domain-containing protein 4C | Q5VZ89 | 0.855917 |
| (C) |  |  |  |
| Gene name | Protein name | Accession | Fold-change |
| H1-4 | Histone H1.4 | P10412 | -1.94501 |
| GSPT1 | Eukaryotic peptide chain release factor subunit 3a | P15170 | -0.97709 |
| H1-2 | Histone H1.2 | P16403 | -0.71206 |
| SUB1 | Activated RNA polymerase II transcriptional coactivator p15 | P53999 | -0.64848 |
| KRT18 | Keratin, type I cytoskeletal 18 | P05783 | -0.56628 |
| ANP32B | Acidic leucine-rich nuclear phosphoprotein 32 family member B | Q92688 | -0.55258 |
| H2AC11 | Histone H2A type 1 | P0C0S8 | -0.53413 |
| EIF3CL | Eukaryotic translation initiation factor 3<br>subunit C-like protein | B5ME19 | -0.42143 |
| TPD52L2 | Tumor protein D54 | O43399 | -0.37034 |
| RPL27A | 60S ribosomal protein L27a | P46776 | -0.36698 |

(D)
| Gene name | Protein name | Accession | Fold-change |
| --- | --- | --- | --- |
| PSMB2 | Proteasome subunit beta type-2 | P49721 | 0.478164 |
| RAB7A | Ras-related protein Rab-7a | P51149 | 0.485954 |
| NAT10 | RNA cytidine acetyltransferase | Q9H0A0 | 0.547291 |
| EIF3E | Eukaryotic translation initiation factor 3<br>subunit E | P60228 | 0.582263 |
| NME1 | Nucleoside diphosphate kinase A | P15531 | 0.600471 |
| RAC2 | Ras-related C3 botulinum toxin substrate 2 | P15153 | 0.61141 |
| EIF3M | Eukaryotic translation initiation factor 3<br>subunit M | Q7L2H7 | 0.616847 |
| COPB1 | Coatomer subunit beta | P53618 | 0.627621 |
| TPD52 | Tumor protein D52 | P55327 | 0.706644 |
| IPO7 | Importin-7 | O95373 | 0.820229 |

## 3. Discussion

Our group previously demonstrated, *in vitro* and *in vivo*, the role of ArtinM in modulating innate [8,22,24,47,48] and adaptive [23,49] immune responses. We have determined that the interaction of ArtinM with N-glycans linked to glycoproteins on the cell surface, such as TLR2/CD14 and CD3εγ, provides the primary mechanism for triggering the production of cytokines responsible for the immunomodulation induced by ArtinM [8,22,50,51]. However, the recognition of TLR2/CD14 N-glycans does not account for the ArtinM activity that induces murine B cells to produce IL-17 and IL-12p40 [26]. Notably, the subcutaneous administration of ArtinM to naive BALB/c mice, besides affecting immune cells, increases the number of splenic B cells [52], suggesting that B cells contribute to the immunomodulatory activity of ArtinM in vivo. To explore this premise, we investigated the effect of ArtinM on isolated murine B cells and found that ArtinM is not directly responsible for B cell proliferation *in vivo*. Instead, lectin promotes murine B cell apoptosis. This cytotoxic effect of ArtinM was also detected in Raji cells, a human B-cell line sourced from NHL. The phosphatase activity of CD45 and protein kinases of the Src family are involved in the cytotoxic effects of ArtinM on Raji cells.

The general characterization of the biological effects of lectins on distinct cell types frequently includes measurement of mitochondrial activity by MTT assay, which allows screening if the explored lectins induce cell proliferation (high mitochondrial activity, likely linked to a cell activation process) or apoptosis (low mitochondrial activity, probably accounting for cytotoxic effects) [53–58]. We found a dose-dependent decrease in mitochondrial activity in B cells that were stimulated for 24 h with ArtinM at the higher concentrations tested, indicating that the lectin could trigger B cell apoptosis. Indeed, it was confirmed that ArtinM diminished the number of viable cells and increased the frequency of double-positive B cells for AnV and PI, which are flow cytometric criteria used to identify cell apoptosis. The association between reduced mitochondrial activity and cytotoxic effects of ArtinM has been reported when stimulating a few cell types, such as murine CD4^+^ and CD8^+^ T cells [23], NB4 human myeloid leukemia cells[27], and Jurkat human leukemic T cells [23]. Beyond the decrease in mitochondrial activity determined by 24 h stimulation with ArtinM at high concentrations, the MTT assay also revealed a small augmentation of mitochondrial activity in B cells stimulated for 48 h with ArtinM at low concentrations compared with the negative control. This could imply the occurrence of B cell proliferation; however, we did not find a concomitant augmentation of thymidine incorporation or IL-2 production by B cells stimulated by ArtinM. Few studies have reported on the effects of lectins on B cell proliferation or apoptosis [4,59]. Interestingly, jacalin, the Galβ1-3Gal-NAc-binding lectin, is also sourced from *A. heterophyllus* seeds and induces B-cell apoptosis, which results from lectin interaction with CD45 O-glycans [60].

We initially aimed to determine the contribution of B cells to the Th1 immune response, which is induced after ArtinM administration to mice experimentally infected with an intracellular pathogen to which they are susceptible. We found that isolated murine B cells stimulated in vitro with ArtinM, even when used at concentrations with low cytotoxic effects, did not respond by producing IL-12 and IFN-γ. Instead, ArtinM-stimulated B cells become apoptotic rather than proinflammatory. Because of the high cytotoxic effect exerted by ArtinM on B cells, we aimed to investigate lectin activity in human B cell lines derived from Burkitt lymphoma, a subtype of NHL [61]. We first confirmed that ArtinM binds to Raji and Daudi cell surface, in a manner dependent on carbohydrate recognition. This phenomenon also occurs with other lectins from different sources having specificity for distinct glycoligands, such as SAL (*Silurus asotus*), which has specificity for α-galactoside ligands [62,63], CGL (*Crenomytilus grayanus*) for Galα1-4Galβ1-4GlcNAc ligands [64], MytiLec (*Mytilus galloprovincialis*) for α-GalNAc ligands [65], and OLL (*Osmerus lanceolatus*) for globotriaosylceramide ligands [66]. By recognizing glycoligands on the Raji and Daudi cell surface, the reported lectins reduce cell viability as ArtinM did. We found that the cytotoxic effect of ArtinM was pronounced in Raji cells, which diminished their growth and augmented their frequency of apoptosis. Diminished Raji cell viability was previously described to be induced by a few lectins, such as jacalin (*Artocarpus heterophyllus*) [67], SAL (*Silurus asotus*) [66], and ALL (*Artocarpus lingnanensis*) [68].

We verified that ArtinM induces Raji cell apoptosis without promoting DNA fragmentation, which is not always required in the apoptosis pathway [69–71]. This is consistent with our previous demonstration that ArtinM induces apoptosis in the human myeloid leukemia cell line (NB4) without causing DNA damage [27]. Abrin, a D-galactose-binding lectin from *Abrus precatorius*, also induces apoptosis in human cell lines derived from myeloid leukemia (MOLT-4) and acute lymphoblastic leukemia (HPB-ALL), which is not associated with DNA fragmentation [72,73]. Our present work showed that ArtinM did not induce Raji cell autophagy, as demonstrated by the reduced levels of relative expression of specific transcripts for the autophagy pathway. In contrast, we previously reported that ArtinM induces autophagy in NB4 cells without caspase involvement [27]. The lectins Abrin (*Abrus precatorius*) and CvL (*Cliona varians*) induce apoptosis in human acute lymphoblastic leukemia (CCRF-CEM) and human erythroleukemia (K562) cell lines, respectively, in a manner that is independent of caspase activity [72,73]. Our study showed that ArtinM-induced apoptosis of Raji cells was not related to the mitochondrial pathway.

To provoke a cell response, extracellular signals activate several signaling pathways to regulate proliferation, survival, cell cycle arrest, and cell death [74]. In this sense, Raji cells apoptosis induced by the lectin ALL, an N-acetyl-D-galactosamine ligand, was reported as being mediated by the p38MAPK signaling pathway [68]. Another study showed that MytiLec, a Galα1-4Galβ1-4Gl specific lectin, recognizes Ramos cells (BL lineage) and induces apoptosis via activation of the MEK/ERK and JNK/p38 MAPK pathways [65]. Various signaling pathways were assessed as being possibly involved in ArtinM-induced Raji cell apoptosis. The pathways were approached by pharmacological inhibition of the protein kinases p38MAPK, JNK, ERK, PKC, and PTK. The role of either molecule was evaluated by measuring the cell growth and frequency of double-positive cells for AnV and PI in B cell cultures after incubation with inhibitors followed by the ArtinM stimulus. The participation of PTK was detected in the ArtinM-stimulated reduction of Raji cell growth, and PTK may be an important negative regulator of ArtinM activity in Raji cells. Our group reported the involvement of PTK in the apoptosis of Jurkat cells (T lymphocyte lines from leukemic cells) induced by ArtinM, which acts as a positive regulator of Jurkat cell activation in response to ArtinM, resulting in the apoptosis of Jurkat cells [23].

To address this hypothesis, the present study also examined the B cell antigen receptor (BCR) signaling cascade as a possible major mediator of Raji cell apoptosis induced by ArtinM. Of the components involved in BCR signaling, the protein kinases of the Src family and CD45 receptor participate strictly in pro-survival signal propagation [61], acting as positive and negative regulators, respectively, of the signaling cascade from BCR [75]. Several studies have reported that CD45 expressed on B cells is targeted by lectins such as PNA (*Arachis hypogaea*) [76], jacalin [60], and Galectin-9 [77]. The recognition of CD45 on B cell surface by Galectin-9 suppresses B cell activation and cell proliferation through Lyn activation, which mediates inhibitory signaling pathways associated with the CD22 co-receptor and SHP-1 tyrosine phosphatase [77]. We investigated the influence of CD45 regulatory activity on the growth and viability of ArtinM-stimulated Raji cells. Using the pharmacological inhibitor NQ301, we blocked the dephosphorylation of the Y505 residue promoted by Lck molecule through CD45 phosphatase activity and verified the enhancement of ArtinM-induced Raji cell apoptosis, suggesting that CD45 phosphatase activity acts as a negative regulator of the lectin effect. It is known that during BCR engagement in chronic lymphoid leukemia (CLL) cells, CD79α phosphorylation is induced by Lck, which induces activation of PI3K/Akt, NF-κB, and MAPK, promoting the survival of stimulated CLL cells [78]. In addition, Src family protein kinases (SFKs), such as Lyn, Fyn, and Blk, are required for several B-cell functions, including differentiation, proliferation, and survival [79]. SFKs are also regulated by CD45 phosphatase activity [80], and their role in Raji cell apoptosis was verified using the pharmacological inhibitor, dasatinib. Inhibition of Src family kinases blocked the ArtinM-stimulated reduction of Raji cell growth and viability. These data indicate that the kinases of the Src family and CD45 phosphatase activity regulate the cytotoxic effect of ArtinM on Raji cells.

The cytotoxic effect of ArtinM was more evident in Raji cells than in Daudi cells. This difference allowed us to establish convenient comparisons of the biological facts determined by ArtinM stimulation in the two cell lines. It has been considered in the proteomic approach, to discriminate discrepant and valuable molecules upregulated or downregulated in each cell line undergoing the ArtinM effects. Our data demonstrated the upregulation of major histocompatibility complex (MHC) class II molecules in ArtinM-stimulated Raji cells. MHC-II molecules, including HLA-DR, HLA-DQ, and HLA-DP, were already reported to participate in the process of inducing cell death [81,82]. Remarkably, the ability of HLA-DR to trigger apoptosis in healthy and neoplastic B-lymphocytes has been reported [83,84]. Consistently, Bertho et al. demonstrated the induction of Raji cell apoptosis via HLA-DR signaling [85]. Apoptosis of B cells triggered by HLA-DR occurs in a caspase-independent manner [86].

Proteomic analysis of ArtinM-stimulated Daudi cells revealed that histones and acidic nuclear phosphoprotein 32 family member B (ANP32B) were the most abundantly detected proteins. Interestingly, ANP32B downregulation exerts anti-apoptotic effects, whereas ANP32B acts as a transcriptional regulator of genes involved in cell cycle progression [87,88]. In human leukemic cell lines, ANP32B downregulation enhances caspase-3 activation, leading to cell apoptosis [89]. ANP32B downregulation has been shown to exert anti-apoptotic effects on hepatocellular carcinoma [90] and breast cancer [91] cells. The ANP32B protein acts as a histone chaperone; its direct binding to histones regulates gene transcription, which affects acetylation and phosphorylation inhibition [92–94]. Taken together, our results suggest that MHC-II molecules are involved in the apoptosis of ArtinM-induced Raji cells, which may explain the absence of DNA fragmentation. However, the regulation triggered by ANP32B in Daudi cells could inhibit the cytotoxic effect of ArtinM in Daudi cells. We conclude that the cytotoxic effect of ArtinM on murine B cells is largely prominent and reproducible in Raji cells, an NHL cell line. Apoptosis induced by ArtinM in B cells derived from NHL is strongly regulated by CD45 phosphatase activity and Src family kinases.

## 4. Materials and Methods

### 4.1. Animals

Male C57BL/6 mice at 6–8 weeks of age were used in this study. The mice were acquired from the animal facility of Ribeirão Preto Medical School at the University of São Paulo, Ribeirão Preto, São Paulo, Brazil. The animals were bred and housed under optimized hygienic conditions at the animal facility of the Department of Cell and Molecular Biology and Pathogenic Bioagents at the Ribeirão Preto Medical School. All experimental procedures were performed in accordance with the local animal ethical committee of Ribeirão Preto Medical School of the University of São Paulo (protocol number: 061/2019).

### 4.2. Purification of ArtinM lectin

ArtinM lectin was purified from the saline extract of *Artocarpus heterophyllus* (jackfruit) seeds using affinity chromatography with D-mannose columns coupled to Sepharose® (Pierce Chemical Company, USA), according to previously described protocols by Da Silva et al. [19].

### 4.3. Murine B cells isolation

Suspensions of spleen cells obtained from mice were prepared as reported by Oliveira-Brito et al. [26] using the B cell enrichment kit (Pan B Cell Isolation Kit, Miltenyi Biotec, EUA), according to the manufacturer’s instructions. The negatively selected cells were stained with PE Rat Anti-Mouse CD19 (BD Pharmingen™) antibodies and analyzed by flow cytometry (Guava® easyCyte, Millipore, EUA). The isolation of B cells reached about 96% of positive cells for CD19 marker.

### 4.4. Burkitt’s Lymphoma cell lines

Raji and Daudi cell lines derived from Burkitt’s lymphoma were routinely grown in advanced Roswell Park Memorial Institute 1640 medium (Gibco®, Life Technologies, Carlsbad, CA, EUA) supplemented with 10% fetal bovine serum, 1% streptomycin/penicillin, 1 mM sodium pyruvate, and 2 mM L-glutamine, in a humidified 5% CO^2^ atmosphere at 37 °C. Raji and Daudi cells were gifts from Dra. Fabíola Traina (Department of Medical Images, Hematology and Clinical Oncology, Hospital of the Ribeirão Preto Medical School, University of São Paulo).

### 4.5. ArtinM binding on Raji and Daudi cell surface

Raji and Daudi cells (1 × 10^6^ cells/mL) were fixed with phosphate-buffered saline (PBS)/3% formaldehyde for 20 min on ice, and washed with PBS or 1% glycine. The cells were then washed twice with PBS and incubated with biotinylated ArtinM (20 µg/mL) that was pre-incubated with or without mannotriose (1 mM), mannose (20 mM), lactose (20 mM), or medium alone for 40 min. The cells were then washed with PBS and incubated with streptavidin-fluorescein isothiocyanate (FITC; 5 µg/mL; Invitrogen) for 40 min. The cells were washed with PBS, and biotinylated ArtinM was detected by flow cytometry (Guava® easyCyte, Millipore, USA). Raji and Daudi cells were incubated with streptavidin-FITC alone (5 µg/mL) in the absence of lectin and were used as a negative control. The percentage of positive cells was plotted as a histogram.

### 4.6. Determination of mitochondrial activity by 3-(4,5-Dimethylthiazol-2-yl)-2,5-Diphenyltetrazolium Bromide (MTT)

Purified murine B cells (5 × 10^5^ cells/well) were stimulated with ArtinM (0.312 - 5.0 µg/mL) for 24 or 48 h at 37 °C. PMA (50 ng/mL) plus ionomycin (1 µM) was used as a positive control for cell activation, and cells incubated with medium alone were used as negative controls. The mitochondrial activity of the cells was determined after the reduction of MTT (Sigma-Aldrich) to produce formazan crystals [95]. The procedure was performed as described by da Silva et al. [49]. Mitochondrial activity was expressed as a percentage after comparison with the absorbance of unstimulated B cells (medium).

### 4.7. Cell proliferation assay and measurement of IL-2 levels

Purified murine B cells (5 × 10^5^ cells/well) were stimulated with ArtinM (0.312 - 5.0 µg/mL), PMA (50 ng/mL) plus ionomycin (1 µM), or the medium alone. After 48 h of incubation at 37 °C, [^3^H]-thymidine ([^3^H]-TdR; Amersham Bioscience, Boston, MA, USA) at 0.5 µCi/well was added and the proliferative rate was measured by [^3^H]-TdR incorporation. The detection of [^3^H]-TdR incorporation was measured using a scintillator, and the results were expressed as counts per minute (CPM) and the stimulation index of cell proliferation. After 48 h of incubation, murine B cells were centrifuged (300 × g, 10 min at room temperature) and the supernatants were collected to measure the levels of IL-2 by ELISA using the OptEIA™ kit (BD Biosciences, USA), according to the manufacturer’s instructions.

### 4.8. Intracellular staining of IL-12 and IFN-γ

Purified murine B cells (5 × 10^5^ cells/well) were stimulated with ArtinM (0.312 - 5.0 µg/mL), PMA (50 ng/mL) plus ionomycin (1 µM), or culture medium alone. After 12 h of incubation at 37 °C, the protein transport inhibitor was added (BD GolgiStopTM; 1 µL every 1.5 mL of the culture), and after 12 h the cells were washed and fixed/permeabilized with BD Cytofix/CytopermTM Plus (20 min at 4 °C). The cells were washed with wash buffer (BD Perm/WashTM buffer) and resuspended in Perm/WashTM buffer with anti-IFN-γ PE (BD Pharmingen™) and anti-IL-12 FITC (BD PharmingenTM) antibodies. After 40 min of incubation at 4 °C, the cells were washed and resuspended in Perm/WashTM buffer, and the frequency of positive B cells for IL-12 and IFN-γ (IL-12^+^/IFN-γ^+^) was measured using flow cytometry (Guava EasyCyte™ Mini System).

### 4.9. Quantification of positive cells for annexin V (AnV) and/or propidium iodide (PI)

Purified murine B cells (5 × 10^5^ cells/well), Raji cells, or Daudi cells (1 × 10^6^ cells/mL) were stimulated with different concentrations of ArtinM (0.019-20.0 µg/mL), PMA (50 ng/mL) plus Ionomycin (1 µM), Arsenic trioxide (AS_2_O_3_; 12 µM or 24 µM) or culture medium alone at 37 °C. After 24 or 48 h, as specified in the figure legends, the cells were stained with annexin V-FITC (5 µg/mL; 40 min at 37 °C), and propidium iodide (10 µg/mL) was added at the time of analysis. The frequency of positive cells for AnV binding and/or PI incorporation was quantified by flow cytometry (Guava EasyCyte™ Mini System).

### 4.10. DNA fragmentation assay

Raji and Daudi cells (1 × 10^6^ cells/mL) were stimulated with ArtinM at concentrations of 1.25 µg/mL and 20.0 µg/mL, respectively, for 24 or 48 h at 37 °C. Arsenic trioxide (AS_2_O_3_) at concentrations of 12 µM (Raji cells) and 24 µM (Daudi cells) was used as a positive control to induce cell death, and the medium alone was used as a negative control. The cells were washed with PBS, lysis buffer was added (50 mM Tris-HCl, 10 mM EDTA, 0.2% Triton-X, 100 μg/mL RNase, and 500 μg/mL proteinase K) and kept under constant agitation for 30 min at 4 °C. Subsequently, DNA fragmentation was evaluated by agarose gel electrophoresis (0.8%), and DNA was stained with SYBR-Safe and visualized by the ChemiDoc gel imaging system (Bio-Rad).

### 4.11. qRT-PCR for determining apoptosis marker expression

Raji cells (1 × 10^6^ cells/mL) were stimulated with ArtinM (1.25 µg/mL), Arsenic trioxide (AS_2_O_3_; 12 µM), or medium alone for 24 h at 37 °C. Total RNA was extracted from Raji cells using TRIzol according to the manufacturer’s instructions, and the RNA was converted into cDNA using the iScript™ cDNA synthesis kit (Bio-Rad). qRT-PCR was performed in a final volume of 10 μL using EvaGreen (Bio-Rad) and the quantification was done by using CFX96 (Bio-Rad) equipment. The cycling conditions were as follows: initial denaturation at 95 °C for 30 s, followed by 40 cycles of denaturation at 95 °C for 5 s, and annealing/extension at 60 °C for 5 s The relative expression of the transcripts was quantified using the ΔΔCt method and normalized to β-actin expression. The specificity of amplification was determined using melting curve analysis. The following PCR primers were used: β-actin (F-GTTGTCGACGACGAGCG, R-GCACAGAGCCTCGCCTT), caspase-3 (F-GAGTCCATTGATTCGCTTCC, R-TCTGGTTTTCGGTGGGTG), Apaf-1 (F-CCTCTCATTTGCTGATGTCG, R-TCACTGCAGATTTTCACCAGA), Smac/Diablo (F-AGCTGAATGTGATTCCTGGC, R-GAAGCTGGAAACCACTTGGA), LC3-I (F-TATCACCGGGATTTTGGTTG, R-GAGAAGACCTTCAAGCAGCG), ATG14 (F-TCTTGCTTGCTCTTAAGTCGG, R-GAGCGGCGATTTCGTCTACT), and ATG-12 (F-CCATCACTGCCAAAACACTC, R-TTGTGGCCTCAGAACAGTTG).

### 4.12. Inhibition assay of signaling molecules in Raji and Daudi cells stimulated with ArtinM

Raji and Daudi cells (1 × 10^6^ cells/mL) were incubated, separately, with the following pharmacological inhibitors: SB202190 (p38MAPK inhibitor), SP600125 (JNK inhibitor), PD98059 (p42/44MAPK inhibitor), H-7 (PKC inhibitor), Genistein (PTK inhibitor), NQ-301 (inhibitor of CD45 phosphatase activity on Lck) and Dasatinib (inhibitor of Lyn, Fyn, and Blk) for 210 minutes at 37 °C. The cells were subsequently stimulated with ArtinM at concentrations of 1.25 µg/mL (Raji cells), and 20.0 µg/mL (Daudi cells), for 24 or 48 h at 37 °C. Arsenic trioxide (AS_2_O_3_) at concentrations of 12 µM (Raji cells) and 24 µM (Daudi cells) was used as a positive control, and the medium alone was used as a negative control. The frequency of positive cells for AnV binding and/or PI incorporation was quantified by flow cytometry (Guava EasyCyte™ Mini System), as described in Section 4.9.

### 4.13. In-solution trypsin digestion

In-solution trypsin digestion was performed according to the protocol described by Pessotti et al. [96]. Briefly, a solution of 6 M guanidine hydrochloride (GuHCl) was added to 100 *μ*g of protein under each condition to a make up a final concentration of 3 M GuHCl, followed by the addition of 5 mM dithiothreitol (DTT) (final concentration). The mixture was incubated at 65 °C for 60 min. Iodoacetamide (IAA) was then added to a final concentration of 15 mM, and the samples were incubated for 60 min at room temperature in the dark. To quench the excess IAA, DTT was added to a final concentration of 15 mM. Clean-up of the samples was performed by the addition of ice-cold acetone (8 volumes) and methanol (1 volume), followed by incubation of the samples for 3 h at -80 °C. After centrifugation at 14,000 × *g* for 10 min, protein pellets were washed twice with one volume of ice-cold methanol and then solubilized with NaOH solution (final concentration of 2.5 mM), followed by the addition of 50 mM HEPES buffer, pH 7.5, to a final volume of 100 mL. Trypsin (Proteomics grade; Sigma, USA) was added at 1:100 (enzyme/substrate) ratio and protein samples were incubated at 37 °C for 18 h. Tryptic peptides were desalted using C18 StageTips [97], dried in a SpeedVac, and re-dissolved in 50 mL of 0.1% formic acid prior to nanoflow liquid chromatography/tandem mass spectrometry (LC-MS/MS) analysis.

### 4.14. Mass spectrometry

LC-MS/MS analyses were performed on a Synapt G2 HDMS mass spectrometer (Waters) coupled to the chromatographic system nanoAcquity UPLC (Waters). Digested peptide samples (7 µg) were loaded onto a trap column (Acquity UPLC M-Class Symmetry C18 Trap Column,100 Å, 5 μm, 300 μm × 25 mm, Waters) at 8 μL/min of loading solution (5% acetonitrile and 0.1% formic acid in water) for 5 min. Then, the mixture of trapped peptides was eluted in an analytical column (Acquity UPLC M-Class HSS T3 Column, 1.8 μm, 300 μm × 150 mm, Waters) with a gradient of 5-35% of phase B (0.1% formic acid in acetonitrile, phase A: 0.1% formic acid in water), over 60 min at a flow rate of 3 µL/min. MS data were acquired in data-independent acquisition (DIA) mode UDMS^E^ using ion mobility separation [98] in the m/z range of 50-2000 and in the resolution mode. Peptide ions were fragmented by collision-induced dissociation (CID), in which collision energies were alternated between low (4 eV) and high energy (ramped from 17 to 60 eV) for precursor and fragment ions, respectively, using scan times of 1.0 s. The ESI source was operated in the positive mode with a capillary voltage of 2.7 kV, block temperature of 100 °C, and cone voltage of 40 V. The column temperature was set to 55 °C. For lock mass correction, a [Glu1]-Fibrinopeptide B solution (500 fmol/mL in 50% methanol, 0.1 % formic acid; Peptide 2.0) was infused through the reference sprayer at 2 μL/min and sampled every 60 s for external calibration [99].

### 4.15. Quantitative Proteomics

Label-free quantification (LFQ) was performed using Progenesis QI for proteomics (Nonlinear Dynamics), as previously reported [100,101]. Briefly, the raw files were loaded into the software, and the peaks were aligned based on the precursor ion retention time of the reference run, which was automatically selected. Default peak-picking parameters were applied. The MS data were processed by the Apex3D module using a low-energy threshold of 750 counts and a high-energy threshold of 50 counts. The MS/MS spectra were exported as a .mgf file for protein identification in PEAKS Studio 7.5 (Bioinformatics Solution, Inc.). A search was performed against *Homo sapiens* sequences in the UniprotKB/Swissprot database (www.uniprot.org; reviewed: 26,577 sequences; downloaded on May 10^th^, 2021). Search parameters were set as follows: mass tolerance of 10 ppm for precursor ions and 0.025 Da for fragment ions, up to two missed cleavage sites allowed for trypsin digestion, maximum false discovery rate (FDR) of 1% at the peptide level, and a minimum of two peptides per protein. Carbamidomethylation of cysteines was selected as a fixed modification, and N-terminal acetylation, methionine oxidation, and asparagine/glutamine deamidation were selected as variable modifications. The identification results were then reimported to the Progenesis QI for proteomics as a .pepXml file. Proteins were quantified as the sum of all unique (non-conflicting) normalized peptide ion abundances corresponding to the protein. Mass spectrometry proteomics data were deposited in the ProteomeXchange Consortium via the PRIDE [1] partner repository with the dataset identifier PXD029399 (Reviewer account details: **Username:**, **Password:** 4TrQpmLp).

### 4.16. Statistical analysis

The results are presented as the mean ± standard deviation (SD). All data were analyzed using Prism 7.0 (GraphPad Software, La Jolla, CA, USA). All statistical determinations for normality were analyzed using the Kolmogorov–Smirnov test with the Dallal–Wilkinson–Lillie test for p-values, and when the distribution could not be assumed to be normal, the Kruskal–Wallis test was used. Statistical determinations of the differences in means between groups were performed using the Kruskal–Wallis test followed by Dunn’s multiple comparison test. The experiments were performed in triplicates or quadruplicates, and three independent assays were performed. Differences with p < 0.05 (*); p < 0.01 (**); p < 0.001 (***); p < 0.0001 (****) were considered statistically significant.

## Funding

This research was financially supported by the Fundação de Amparo a Pesquisa do Estado de São Paulo (grant numbers 2019/05867-9, 2018/21708-5, 2018/18538-0, 2017/20106-9, 2016/04877-2), the Coordenação de Aperfeiçoamento de Pessoal de Nível Superior, CNPq.

## Acknowledgements

We thank Patrícia Vendruscolo, and Érica Vendruscolo for technical support.. We thank Prof. Dr. Reinaldo Salomão from UNIFESP for the use of Progenesis QI for proteomics.

## Author contributions

Conceptualization and designed the study – B.R.B., M.C.R.-B., and T.A.d.S.; planned the experiments and analyzed the data – B.R.B., A.K.T., A.Z., and T.A.d.S.; performed the experiments – B.R.B., S.M.O.T., A.d.C.J., E.M.E., J.G.M., A.K.T., M.F.C., A.Z., and T.A.d.S.; writing/preparation original draf – B.R.B., and T.A.d.S.; review and editing – M.C.R.-B and T.A.d.S; supervision – T.A.d.S. All authors have read and agreed to the published version of the manuscript.

## Data availability statement

All data presented this study are available from the corresponding author, upon responsible request.

## Conflicts of Interest

The authors declare no conflict of interest.

## Notes

### Competing Interest Statement

The authors have declared no competing interest.

## References

1. Mazola, Y.; Chinea, G.; Musacchio, A. Glycosylation and bioinformatics: Current status for glycosylation prediction tools. Biotecnol. Apl. 2011.

2. Pinho, S.S.; Reis, C.A. Glycosylation in cancer: Mechanisms and clinical implications. Nat. Rev. Cancer 2015.

3. Smith, B.A.H.; Bertozzi, C.R. The clinical impact of glycobiology: targeting selectins, Siglecs and mammalian glycans. Nat. Rev. Drug Discov. 2021, 20, 217–243, doi:10.1038/s41573-020-00093-1.

4. Wilhelm, I.; Levit-Zerdoun, E.; Jakob, J.; Villringer, S.; Frensch, M.; Übelhart, R.; Landi, A.; Müller, P.; Imberty, A.; Thuenauer, R.; et al. Carbohydrate-dependent B cell activation by fucose-binding bacterial lectins. Sci. Signal. 2019, doi:10.1126/scisignal.aao7194.

5. Ricci-Azevedo, R.; Roque-Barreira, M.-C.; Gay, N.J. Targeting and Recognition of Toll-Like Receptors by Plant and Pathogen Lectins. Front. Immunol. 2017, 8, 1820, doi:10.3389/fimmu.2017.01820.

6. Rheinländer, A.; Schraven, B.; Bommhardt, U. CD45 in human physiology and clinical medicine. Immunol. Lett. 2018.

7. Shatnyeva, O.M.; Kubarenko, A. V.; Weber, C.E.M.; Pappa, A.; Schwartz-Albiez, R.; Weber, A.N.R.; Krammer, P.H.; Lavrik, I.N. Modulation of the CD95-induced apoptosis: The role of CD95 N-glycosylation. PLoS One 2011, doi:10.1371/journal.pone.0019927.

8. Da Silva, T.A.; Zorzetto-Fernandes, A.L.V.; Cecílio, N.T.; Sardinha-Silva, A.; Fernandes, F.F.; Roque-Barreira, M.C. CD14 is critical for TLR2-mediated M1 macrophage activation triggered by N-glycan recognition. Sci. Rep. 2017, 7, 1–14, doi:10.1038/s41598-017-07397-0.

9. Van Damme, E.J.M.; Peumans, W.J.; Barre, A.; Rougé, P. Plant lectins: A composite of several distinct families of structurally and evolutionary related proteins with diverse biological roles. CRC. Crit. Rev. Plant Sci. 1998, 17, 575–692, doi:10.1016/S0735-2689(98)00365-7.

10. Van Damme, E.J.M.; Lannoo, N.; Peumans, W.J. Plant lectins. Adv. Bot. Res. 2008, 48, 107–209.

11. Carvalho, E.V.M.M.; Oliveira, W.F.; Coelho, L.C.B.B.; Correia, M.T.S. Lectins as mitosis stimulating factors: Briefly reviewed. Life Sci. 2018, 207, 152–157, doi:10.1016/j.lfs.2018.06.003.

12. Schnaar, R.L. Glycans and glycan-binding proteins in immune regulation: A concise introduction to glycobiology for the allergist. J. Allergy Clin. Immunol. 2015.

13. Lichtenstein, R.G.; Rabinovich, G.A. Glycobiology of cell death: When glycans and lectins govern cell fate. Cell Death Differ. 2013.

14. Peumans, W.J.; Van Damme, E.J. Lectins as plant defense proteins. Plant Physiol. 1995.

15. Sharon, N.; Lis, H. History of lectins: From hemagglutinins to biological recognition molecules. Glycobiology 2004.

16. Tsaneva, M.; Van Damme, E.J.M. 130 years of Plant Lectin Research. Glycoconj. J. 2020, doi:10.1007/s10719-020-09942-y.

17. Jandú, J.J.B.; Moraes Neto, R.N.; Zagmignan, A.; de Sousa, E.M.; Brelaz-de-Castro, M.C.A.; dos Santos Correia, M.T.; da Silva, L.C.N. Targeting the immune system with plant lectins to combat microbial infections. Front. Pharmacol. 2017, 8, 1–11, doi:10.3389/fphar.2017.00671.

18. Johannssen, T.; Lepenies, B. Glycan-Based Cell Targeting To Modulate Immune Responses. Trends Biotechnol. 2017, 35, 334–346, doi:10.1016/j.tibtech.2016.10.002.

19. da Silva, T.A.; Oliveira-Brito, P.K.M.; de Oliveira Thomaz, S.M.; Roque-Barreira, M.C. ArtinM: Purification and Evaluation of Biological Activities. In Methods in Molecular Biology; 2020; pp. 349–358.

20. Santos-de-oliveira, R.; Dias-baruffi, M.; Thomaz, S.M.; Beltrarnini, L.M.; Roque-barreira, M.C. A neutrophil migration-inducing lectin from Artocarpus integrifolia. J. Immunol. 1994, 153, 1798–1897.

21. Liu, Y.; Cecílio, N.T.; Carvalho, F.C.; Roque-Barreira, M.C.; Feizi, T. Glycan microarray analysis of the carbohydrate-recognition specificity of native and recombinant forms of the lectin ArtinM. Data Br. 2015, 5, 1035–1047, doi:10.1016/j.dib.2015.11.014.

22. Mariano, V.S.; Zorzetto-Fernandes, A.L.; Da Silva, T.A.; Ruas, L.P.; Nohara, L.L.; De Almeida, I.C.; Roque-Barreira, M.C. Recognition of TLR2 N-glycans: Critical role in ArtinM immunomodulatory activity. PLoS One 2014, 9, 1–9, doi:10.1371/journal.pone.0098512.

23. da Silva, T.A.; Oliveira-Brito, P.K.M.; Gonçalves, T.E.; Vendruscolo, P.E.; Roque-Barreira, M.C. Artinm mediates murine T cell activation and induces cell death in jurkat human leukemic T cells. Int. J. Mol. Sci. 2017, 18, 1–21, doi:10.3390/ijms18071400.

24. Barbosa-Lorenzi, V.C.; Buranello, P.A. de A.; Roque-Barreira, M.C.; Jamur, M.C.; Oliver, C.; Pereira-da-Silva, G. The lectin ArtinM binds to mast cells inducing cell activation and mediator release. Biochem. Biophys. Res. Commun. 2011, 416, 318–324, doi:10.1016/j.bbrc.2011.11.033.

25. Pereira-Da-Silva, G.; Moreno, A.N.; Marques, F.; Oliver, C.; Célia Jamur, M.; Panunto-Castelo, A.; Roque-Barreira, M.C. Neutrophil activation induced by the lectin KM+ involves binding to CXCR2. Biochim. Biophys. Acta - Gen. Subj. 2006, 1760, 86–94, doi:10.1016/j.bbagen.2005.09.011.

26. Oliveira-Brito, P.K.M.; Roque-Barreira, M.C.; Da Silva, T.A. The Response of IL-17-Producing B Cells to ArtinM is independent of its interaction with TLR2 and CD14. Molecules 2018, 23, 1–8.

27. Carvalho, F.C.; Soares, S.G.; Tamarozzi, M.B.; Rego, E.M.; Roque-Barreira, M.C. The recognition of N-glycans by the lectin ArtinM mediates cell death of a human myeloid leukemia cell line. PLoS One 2011, doi:10.1371/journal.pone.0027892.

28. Montecino-Rodriguez, E.; Dorshkind, K. B-1 B Cell Development in the Fetus and Adult. Immunity 2012.

29. Ghia, P.; ten Boekel, E.; Rolink, A.G.; Melchers, F. B-cell development: a comparison between mouse and man. Immunol. Today 1998, 19, 480–485, doi:10.1016/S0167-5699(98)01330-9.

30. Eibel, H.; Kraus, H.; Sic, H.; Kienzler, A.-K.; Rizzi, M. B cell Biology: An Overview. Curr. Allergy Asthma Rep. 2014, 14, 434, doi:10.1007/s11882-014-0434-8.

31. Pieper, K; Grimbacher, B; Eibel, H. B-cell biology and development. J Allergy Clin Immunol 2013, 131, 959–971, doi:10.1016/j.jaci.2013.01.046.

32. Schweighoffer, E.; Tybulewicz, V.L. Signalling for B cell survival. Curr. Opin. Cell Biol. 2018, 8–14.

33. Popi, A.F.; Longo-Maugéri, I.M.; Mariano, M. An overview of B-1 cells as antigen-presenting cells. Front. Immunol. 2016.

34. Davis, R.S. FCRL regulation in innate-like B cells. Ann. N. Y. Acad. Sci. 2015, doi:10.1111/nyas.12771.

35. Aziz, M.; Holodick, N.E.; Rothstein, T.L.; Wang, P. The role of B-1 cells in inflammation. Immunol. Res. 2015.

36. Cerutti, A.; Cols, M.; Puga, I. Marginal zone B cells: Virtues of innate-like antibody-producing lymphocytes. Nat. Rev. Immunol. 2013.

37. Genestier, L.; Taillardet, M.; Mondiere, P.; Gheit, H.; Bella, C.; Defrance, T. TLR Agonists Selectively Promote Terminal Plasma Cell Differentiation of B Cell Subsets Specialized in Thymus-Independent Responses. J. Immunol. 2012, 178, 7779–7786, doi:10.4049/jimmunol.178.12.7779.

38. Hoek, K.L.; Gordy, L.E.; Collins, P.L.; Parekh, V. V.; Aune, T.M.; Joyce, S.; Thomas, J.W.; Van Kaer, L.; Sebzda, E. Follicular B Cell Trafficking within the Spleen Actively Restricts Humoral Immune Responses. Immunity 2010, doi:10.1016/j.immuni.2010.07.016.

39. Mesin, L.; Ersching, J.; Victora, G.D. Germinal Center B Cell Dynamics. Immunity 2016.

40. Shaffer, A.L.; Young, R.M.; Staudt, L.M. Pathogenesis of human B cell lymphomas. Annu. Rev. Immunol. 2012.

41. Fuertes, T.; Ramiro, A.R.; de Yebenes, V.G. miRNA-Based Therapies in B Cell Non-Hodgkin Lymphoma. Trends Immunol. 2020, 41, 932–947, doi:10.1016/j.it.2020.08.006.

42. Mlynarczyk, C.; Fontán, L.; Melnick, A. Germinal center-derived lymphomas: The darkest side of humoral immunity. Immunol. Rev. 2019.

43. Saleh, K.; Michot, J.M.; Camara-Clayette, V.; Vassetsky, Y.; Ribrag, V. Burkitt and Burkitt-Like Lymphomas: a Systematic Review. Curr. Oncol. Rep. 2020.

44. Robaina, M.C.; Mazzoccoli, L.; Klumb, C.E. Germinal Centre B Cell Functions and Lymphomagenesis: Circuits Involving MYC and MicroRNAs. Cells 2019.

45. Asadbeigi, S.N.; Deel, C.D. Burkitt-Like Lymphoma with 11q Aberration: A Case Report and Review of a Rare Entity. Case Rep. Hematol. 2020, 2020, 1–6, doi:10.1155/2020/8896322.

46. Zhang, J.; Meng, L.; Jiang, W.; Zhang, H.; Zhou, A.; Zeng, N. Identification of clinical molecular targets for childhood Burkitt lymphoma. Transl. Oncol. 2020, 13, 100855, doi:10.1016/j.tranon.2020.100855.

47. Toledo, K.A.; Scwartz, C.; Oliveira, A.F.; Conrado, M.C.A.V.; Bernardes, E.S.; Fernandes, L.C.; Roque-Barreira, M.C.; Pereira-da-Silva, G.; Moreno, A.N. Neutrophil activation induced by ArtinM: Release of inflammatory mediators and enhancement of effector functions. Immunol. Lett. 2009, 123, 14–20, doi:10.1016/j.imlet.2009.01.009.

48. Ricci-Azevedo, R.; Oliveira, A.F.; Conrado, M.C.A.V.; Carvalho, F.C.; Roque-Barreira, M.C. Neutrophils Contribute to the Protection Conferred by ArtinM against Intracellular Pathogens: A Study on Leishmania major. PLoS Negl. Trop. Dis. 2016, 10, 1–23, doi:10.1371/journal.pntd.0004609.

49. Da Silva, T.A.; De Souza, M.A.; Cecílio, N.T.; Roque-Barreira, M.C. Activation of spleen cells by ArtinM may account for its immunomodulatory properties. Cell Tissue Res. 2014, 357, 719–730, doi:10.1007/s00441-014-1879-8.

50. Teixeira, C.R.; Cavassani, K.A.; Gomes, R.B.; Teixeira, M.J.; Roque-Barreira, M.C.; Cavada, B.S.; Silva, J.S. Da; Barral, A.; Barral-Netto, M. Potential of KM+ lectin in immunization against Leishmania amazonensis infection. Vaccine 2006, 24, 3001–3008, doi:10.1016/j.vaccine.2005.11.067.

51. Souza, M.A.; Carvalho, F.C.; Ruas, L.P.; Ricci-Azevedo, R.; Roque-Barreira, M.C. The immunomodulatory effect of plant lectins: A review with emphasis on ArtinM properties. Glycoconj. J. 2013, 30, 641–657, doi:10.1007/s10719-012-9464-4.

52. Oliveira Brito, P.K.M.; Gonçalves, T.E.; Fernandes, F.F.; Miguel, C.B.; Rodrigues, W.F.; Lazo Chica, J.E.; Roque-Barreira, M.C.; Da Silva, T.A. Systemic effects in naïve mice injected with immunomodulatory lectin ArtinM. PLoS One 2017, 12, 1–19, doi:10.1371/journal.pone.0187151.

53. Nolte, S.; de Castro Damasio, D.; Baréa, A.C.; Gomes, J.; Magalhães, A.; Mello Zischler, L.F.C.; Stuelp-Campelo, P.M.; Elífio-Esposito, S.L.; Roque-Barreira, M.C.; Reis, C.A.; et al. BJcuL, a lectin purified from Bothrops jararacussu venom, induces apoptosis in human gastric carcinoma cells accompanied by inhibition of cell adhesion and actin cytoskeleton disassembly. Toxicon 2012, 59, 81–85, doi:10.1016/j.toxicon.2011.10.012.

54. Casella-Martins, A.; Ayres, L.R.; Burin, S.M.; Morais, F.R.; Pereira, J.C.; Faccioli, L.H.; Sampaio, S. V.; Arantes, E.C.; Castro, F.A.; Pereira-Crott, L.S. Immunomodulatory activity of Tityus serrulatus scorpion venom on human T lymphocytes. J. Venom. Anim. Toxins Incl. Trop. Dis. 2015, doi:10.1186/s40409-015-0046-3.

55. Palharini, J.G.; Richter, A.C.; Silva, M.F.; Ferreira, F.B.; Pirovani, C.P.; Naves, K.S.C.; Goulart, V.A.; Mineo, T.W.P.; Silva, M.J.B.; Santiago, F.M. Eutirucallin: A lectin with antitumor and antimicrobial properties. Front. Cell. Infect. Microbiol. 2017, doi:10.3389/fcimb.2017.00136.

56. da Silva, P.M.; de Moura, M.C.; Gomes, F.S.; da Silva Trentin, D.; Silva de Oliveira, A.P.; de Mello, G.S.V.; da Rocha Pitta, M.G.; de Melo Rego, M.J.B.; Coelho, L.C.B.B.; Macedo, A.J.; et al. PgTeL, the lectin found in Punica granatum juice, is an antifungal agent against Candida albicans and Candida krusei. Int. J. Biol. Macromol. 2018, doi:10.1016/j.ijbiomac.2017.12.039.

57. da Silva, J.D.F.; da Silva, S.P.; da Silva, P.M.; Vieira, A.M.; de Araújo, L.C.C.; de Albuquerque Lima, T.; de Oliveira, A.P.S.; do Nascimento Carvalho, L.V.; da Rocha Pitta, M.G.; de Melo Rêgo, M.J.B.; et al. Portulaca elatior root contains a trehalose-binding lectin with antibacterial and antifungal activities. Int. J. Biol. Macromol. 2019, doi:10.1016/j.ijbiomac.2018.12.188.

58. Procópio, T.F.; de Siqueira Patriota, L.L.; de Moura, M.C.; da Silva, P.M.; de Oliveira, A.P.S.; do Nascimento Carvalho, L.V.; de Albuquerque Lima, T.; Soares, T.; da Silva, T.D.; Breitenbach Barroso Coelho, L.C.; et al. CasuL: A new lectin isolated from Calliandra surinamensis leaf pinnulae with cytotoxicity to cancer cells, antimicrobial activity and antibiofilm effect. Int. J. Biol. Macromol. 2017, 98, 419–429, doi:10.1016/j.ijbiomac.2017.02.019.

59. Bhutia, S.K.; Mallick, S.K.; Maiti, T.K. In vitro immunostimulatory properties of Abrus lectins derived peptides in tumor bearing mice. Phytomedicine 2009, doi:10.1016/j.phymed.2009.01.006.

60. Ma, B.Y.; Yoshida, K.; Baba, M.; Nonaka, M.; Matsumoto, S.; Kawasaki, N.; Asano, S.; Kawasaki, T. The lectin Jacalin induces human B-lymphocyte apoptosis through glycosylation-dependent interaction with CD45. Immunology 2009, doi:10.1111/j.1365-2567.2008.02977.x.

61. Valla, K.; Flowers, C.R.; Koff, J.L. Targeting the B cell receptor pathway in non-Hodgkin lymphoma. Expert Opin. Investig. Drugs 2018, 27, 513–522, doi:10.1080/13543784.2018.1482273.

62. Sugawara, S.; Kawano, T.; Omoto, T.; Hosono, M.; Tatsuta, T.; Nitta, K. Binding of Silurus asotus lectin to Gb3 on Raji cells causes disappearance of membrane-bound form of HSP70. Biochim. Biophys. Acta -Gen. Subj. 2009, doi:10.1016/j.bbagen.2008.10.005.

63. Kawano, T.; Sugawara, S.; Hosono, M.; Tatsuta, T.; Ogawa, Y.; Fujimura, T.; Taka, H.; Murayama, K.; Nitta, K. Globotriaosylceramide-expressing Burkitt’s lymphoma cells are committed to early apoptotic status by rhamnose-binding lectin from catfish eggs. Biol. Pharm. Bull. 2009, doi:10.1248/bpb.32.345.

64. Chernikov, O.; Kuzmich, A.; Chikalovets, I.; Molchanova, V.; Hua, K.F. Lectin CGL from the sea mussel Crenomytilus grayanus induces Burkitt’s lymphoma cells death via interaction with surface glycan. Int. J. Biol. Macromol. 2017, doi:10.1016/j.ijbiomac.2017.06.074.

65. Hasan, I.; Sugawara, S.; Fujii, Y.; Koide, Y.; Terada, D.; Iimura, N.; Fujiwara, T.; Takahashi, K.G.; Kojima, N.; Rajia, S.; et al. Mytilec, a mussel R-type lectin, interacts with surface glycan GB3 on burkitt’s lymphoma cells to trigger apoptosis through multiple pathways. Mar. Drugs 2015, doi:10.3390/md13127071.

66. Hosono, M.; Sugawara, S.; Matsuda, A.; Tatsuta, T.; Koide, Y.; Hasan, I.; Ozeki, Y.; Nitta, K. Binding profiles and cytokine-inducing effects of fish rhamnose-binding lectins on Burkitt’s lymphoma Raji cells. Fish Physiol. Biochem. 2014, doi:10.1007/s10695-014-9948-1.

67. Baba, M.; Yong Ma, B.; Nonaka, M.; Matsuishi, Y.; Hirano, M.; Nakamura, N.; Kawasaki, N.; Kawasaki, N.; Kawasaki, T. Glycosylation-dependent interaction of Jacalin with CD45 induces T lymphocyte activation and Th1/Th2 cytokine secretion. J. Leukoc. Biol. 2007, doi:10.1189/jlb.1106660.

68. Luo, Y.; Liu, X.; Lin, F.; Liao, L.; Deng, Y.; Zeng, L.; Zeng, Q. Cloning of a novel lectin from artocarpus lingnanensis that induces apoptosis in human B-lymphoma cells. Biosci. Biotechnol. Biochem. 2018, doi:10.1080/09168451.2017.1415127.

69. Schulze-Osthoff, K.; Walczak, H.; Dröge, W.; Krammer, P.H. Cell nucleus and DNA fragmentation are not required for apoptosis. J. Cell Biol. 1994, doi:10.1083/jcb.127.1.15.

70. Zhang, J.; Liu, X.; Scherer, D.C.; Van Kaer, L.; Wang, X.; Xu, M. Resistance to DNA fragmentation and chromatin condensation in mice lacking the DNA fragmentation factor 45. Proc. Natl. Acad. Sci. U. S. A. 1998, doi:10.1073/pnas.95.21.12480.

71. Zhang, J.H.; Xu, M. DNA fragmentation in apoptosis. Cell Res. 2000.

72. Queiroz, A.F.S.; Silva, R.A.; Moura, R.M.; Dreyfuss, J.L.; Paredes-Gamero, E.J.; Souza, A.C.S.; Tersariol, I.L.S.; Santos, E.A.; Nader, H.B.; Justo, G.Z.; et al. Growth inhibitory activity of a novel lectin from Cliona varians against K562 human erythroleukemia cells. Cancer Chemother. Pharmacol. 2009, doi:10.1007/s00280-008-0825-4.

73. Ohba, H.; Moriwaki, S.; Bakalova, R.; Yasuda, S.; Yamasaki, N. Plant-derived abrin-a induces apoptosis in cultured leukemic cell lines by different mechanisms. Toxicol. Appl. Pharmacol. 2004, doi:10.1016/j.taap.2003.11.018.

74. Wada, T.; Penninger, J.M. Mitogen-activated protein kinases in apoptosis regulation. Oncogene 2004.

75. Zikherman, J.; Parameswaran, R.; Hermiston, M.; Weiss, A. The Structural Wedge Domain of the Receptor-like Tyrosine Phosphatase CD45 Enforces B Cell Tolerance by Regulating Substrate Specificity. J. Immunol. 2013, doi:10.4049/jimmunol.1202928.

76. Giovannone, N.; Antonopoulos, A.; Liang, J.; Geddes Sweeney, J.; Kudelka, M.R.; King, S.L.; Lee, G.S.; Cummings, R.D.; Dell, A.; Barthel, S.R.; et al. Human B Cell Differentiation Is Characterized by Progressive Remodeling of O-Linked Glycans. Front. Immunol. 2018, doi:10.3389/fimmu.2018.02857.

77. Giovannone, N.; Liang, J.; Antonopoulos, A.; Geddes Sweeney, J.; King, S.L.; Pochebit, S.M.; Bhattacharyya, N.; Lee, G.S.; Dell, A.; Widlund, H.R.; et al. Galectin-9 suppresses B cell receptor signaling and is regulated by I-branching of N-glycans. Nat. Commun. 2018, doi:10.1038/s41467-018-05770-9.

78. Talab, F.; Allen, J.C.; Thompson, V.; Lin, K.; Slupsky, J.R. LCK is an important mediator of B-cell receptor signaling in chronic lymphocytic leukemia cells. Mol. Cancer Res. 2013, doi:10.1158/1541-7786.MCR-12-0415-T.

79. Roskoski, R. Src kinase regulation by phosphorylation and dephosphorylation. Biochem. Biophys. Res. Commun. 2005.

80. Zikherman, J.; Doan, K.; Parameswaran, R.; Raschke, W.; Weiss, A. Quantitative differences in CD45 expression unmask functions for CD45 in B-cell development, tolerance, and survival. Proc. Natl. Acad. Sci. U. S. A. 2012, doi:10.1073/pnas.1117374108.

81. Bertho, N.; Drénou, B.; Laupeze, B.; Berre, C. Le; Amiot, L.; Grosset, J.-M.; Fardel, O.; Charron, D.; Mooney, N.; Fauchet, R. HLA-DR-Mediated Apoptosis Susceptibility Discriminates Differentiation Stages of Dendritic/Monocytic APC. J. Immunol. 2000, 164, 2379–2385, doi:10.4049/jimmunol.164.5.2379.

82. Bertho, N. MHC class II-mediated apoptosis of mature dendritic cells proceeds by activation of the protein kinase C-delta isoenzyme. Int. Immunol. 2002, 14, 935–942, doi:10.1093/intimm/dxf058.

83. Truman, J.-P.; Ericson, M.L.; Choqueux-Séébold, C.J.M.; Charron, D.J.; Mooney, N.A. Lymphocyte programmed cell death is mediated via HLA class II DR. Int. Immunol. 1994, 6, 887–896, doi:10.1093/intimm/6.6.887.

84. Nagy, Z.A.; Hubner, B.; Löhning, C.; Rauchenberger, R.; Reiffert, S.; Thomassen-Wolf, E.; Zahn, S.; Leyer, S.; Schier, E.M.; Zahradnik, A.; et al. Fully human, HLA-DR-specific monoclonal antibodies efficiently induce programmed death of malignant lymphoid cells. Nat. Med. 2002, 8, 801–807, doi:10.1038/nm736.

85. Bertho, N.; Laupèze, B.; Mooney, N.; Le Berre, C.; Charron, D.; Drénou, B.; Fauchet, R. HLA-DR mediated cell death is associated with, but not induced by TNF-α secretion in APC. Hum. Immunol. 2001, 62, 106–112, doi:10.1016/S0198-8859(00)00240-8.

86. Guo, W.; Castaigne, J.-G.; Mooney, N.; Charron, D.; Al-Daccak, R. Signaling through HLA-DR induces PKCβ-dependent B cell death outside rafts. Eur. J. Immunol. 2003, 33, 928–938, doi:10.1002/eji.200323351.

87. Sun, W.; Kimura, H.; Hattori, N.; Tanaka, S.; Matsuyama, S.; Shiota, K. Proliferation related acidic leucine-rich protein PAL31 functions as a caspase-3 inhibitor. Biochem. Biophys. Res. Commun. 2006, 342, 817–823, doi:10.1016/j.bbrc.2006.02.026.

88. Sun, W.; Hattori, N.; Mutai, H.; Toyoshima, Y.; Kimura, H.; Tanaka, S.; Shiota, K. PAL31, a Nuclear Protein Required for Progression to the S Phase. Biochem. Biophys. Res. Commun. 2001, 280, 1048–1054, doi:10.1006/bbrc.2000.4244.

89. Shen, S.-M.; Yu, Y.; Wu, Y.-L.; Cheng, J.-K.; Wang, L.-S.; Chen, G.-Q. Downregulation of ANP32B, a novel substrate of caspase-3, enhances caspase-3 activation and apoptosis induction in myeloid leukemic cells. Carcinogenesis 2010, 31, 419–426, doi:10.1093/carcin/bgp320.

90. Ohno, Y.; Koizumi, M.; Nakayama, H.; Watanabe, T.; Hirooka, M.; Tokumoto, Y.; Kuroda, T.; Abe, M.; Fukuda, S.; Higashiyama, S.; et al. Downregulation of ANP32B exerts anti-apoptotic effects in hepatocellular carcinoma. PLoS One 2017, 12, e0177343, doi:10.1371/journal.pone.0177343.

91. Yang, S.; Zhou, L.; Reilly, P.T.; Shen, S.-M.; He, P.; Zhu, X.-N.; Li, C.-X.; Wang, L.-S.; Mak, T.W.; Chen, G.-Q.; et al. ANP32B deficiency impairs proliferation and suppresses tumor progression by regulating AKT phosphorylation. Cell Death Dis. 2016, 7, e2082, doi:10.1038/cddis.2016.8.

92. Munemasa, Y.; Suzuki, T.; Aizawa, K.; Miyamoto, S.; Imai, Y.; Matsumura, T.; Horikoshi, M.; Nagai, R. Promoter region-specific histone incorporation by the novel histone chaperone ANP32B and DNA-binding factor KLF5. Mol. Cell. Biol. 2008, 28, 1171–81, doi:10.1128/MCB.01396-07.

93. Chemnitz, J.; Pieper, D.; Stich, L.; Schumacher, U.; Balabanov, S.; Spohn, M.; Grundhoff, A.; Steinkasserer, A.; Hauber, J.; Zinser, E. The acidic protein rich in leucines Anp32b is an immunomodulator of inflammation in mice. Sci. Rep. 2019, 9, 4853, doi:10.1038/s41598-019-41269-z.

94. Tochio, N.; Umehara, T.; Munemasa, Y.; Suzuki, T.; Sato, S.; Tsuda, K.; Koshiba, S.; Kigawa, T.; Nagai, R.; Yokoyama, S. Solution structure of histone chaperone ANP32B: interaction with core histones H3-H4 through its acidic concave domain. J. Mol. Biol. 2010, 401, 97–114, doi:10.1016/j.jmb.2010.06.005.

95. Mosmann, T. Rapid colorimetric assay for cellular growth and survival: Application to proliferation and cytotoxicity assays. J. Immunol. Methods 1983, 65, 55–63, doi:10.1016/0022-1759(83)90303-4.

96. Pessotti, D.S.; Andrade-Silva, D.; Serrano, S.M.T.; Zelanis, A. Heterotypic signaling between dermal fibroblasts and melanoma cells induces phenotypic plasticity and proteome rearrangement in malignant cells. Biochim. Biophys. Acta - Proteins Proteomics 2020, 1868, 140525, doi:10.1016/j.bbapap.2020.140525.

97. Rappsilber, J.; Mann, M.; Ishihama, Y. Protocol for micro-purification, enrichment, pre-fractionation and storage of peptides for proteomics using StageTips. Nat. Protoc. 2007, 2, 1896–1906, doi:10.1038/nprot.2007.261.

98. Distler, U.; Kuharev, J.; Navarro, P.; Levin, Y.; Schild, H.; Tenzer, S. Drift time-specific collision energies enable deep-coverage data-independent acquisition proteomics. Nat. Methods 2014, 11, 167–170, doi:10.1038/nmeth.2767.

99. Pedroso, A.P.; Souza, A.P.; Dornellas, A.P.S.; Oyama, L.M.; Nascimento, C.M.O.; Santos, G.M.S.; Rosa, J.C.; Bertolla, R.P.; Klawitter, J.; Christians, U.; et al. Intrauterine Growth Restriction Programs the Hypothalamus of Adult Male Rats: Integrated Analysis of Proteomic and Metabolomic Data. J. Proteome Res. 2017, 16, 1515–1525, doi:10.1021/acs.jproteome.6b00923.

100. Câmara, G.A.; Nishiyama-Jr, M.Y.; Kitano, E.S.; Oliveira, U.C.; Silva, P.I. da; Junqueira-de-Azevedo, I.L.; Tashima, A.K. A Multiomics Approach Unravels New Toxins With Possible In Silico Antimicrobial, Antiviral, and Antitumoral Activities in the Venom of Acanthoscurria rondoniae. Front. Pharmacol. 2020, 11, doi:10.3389/fphar.2020.01075.

101. Simizo, A.; Kitano, E.S.; Sant’Anna, S.S.; Grego, K.F.; Tanaka-Azevedo, A.M.; Tashima, A.K. Comparative gender peptidomics of Bothrops atrox venoms: are there differences between them? J. Venom. Anim. Toxins Incl. Trop. Dis. 2020, 26, e20200055, doi:10.1590/1678-9199-JVATITD-2020-0055.

